# Preventing evolutionary rescue in cancer

**DOI:** 10.1101/2023.11.22.568336

**Authors:** Srishti Patil, Armaan Ahmed, Yannick Viossat, Robert Noble

**Affiliations:** Indian Institute of Science Education and Research, Pune, India; Department of Mathematics, City, University of London, London, UK; Department of Applied Math & Statistics, Johns Hopkins University, Baltimore, USA; Department of Biophysics, Johns Hopkins University, Baltimore, USA; Ceremade, CNRS, Université Paris-Dauphine, Université PSL, Paris, France

**Keywords:** mathematical oncology, evolutionary therapy, evolutionary rescue, therapeutic resistance, cancer treatment

## Abstract

First-line cancer treatment frequently fails due to initially rare therapeutic resistance. An important clinical question is then how to schedule subsequent treatments to maximize the probability of tumour eradication. Here, we provide a theoretical solution to this problem by using mathematical analysis and extensive stochastic simulations within the framework of evolutionary rescue theory to determine how best to exploit the vulnerability of small tumours to stochastic extinction. Whereas standard clinical practice is to wait for evidence of relapse, we confirm a recent hypothesis that the optimal time to switch to a second treatment is when the tumour is close to its minimum size before relapse, when it is likely undetectable. This optimum can lie slightly before or slightly after the nadir, depending on tumour parameters. Given that this exact time point may be difficult to determine in practice, we study windows of high extinction probability that lie around the optimal switching point, showing that switching after the relapse has begun is typically better than switching too early. We further reveal how treatment dose and tumour demographic and evolutionary parameters influence the predicted clinical outcome, and we determine how best to schedule drugs of unequal efficacy. Our work establishes a foundation for further experimental and clinical investigation of this evolutionarily-informed “extinction therapy” strategy.

## 1 Introduction

Just as species in an ecosystem interact, compete for resources, adapt to changing environmental conditions and undergo natural selection, so cancer clones rise and fall in a tumour ecosystem. Darwinian principles inevitably determine therapeutic responses [1] including the emergence of resistance, which, despite pharmaceutical advances, remains the greatest challenge in oncology. As cancer cells can use a variety of adaptive strategies to achieve resistance [2], targeting a single molecular mechanism often proves ineffective in the long term [3]. Understanding intratumour evolutionary processes provides a rational foundation for developing treatment strategies that, by explicitly accounting for evolutionary dynamics, achieve better clinical outcomes [4, 5, 6]. Mathematical modelling of clonal dynamics and the emergence of resistance is critical for optimising clinical treatment strategies based on evolutionary principles. Consequently, the historical development of evolutionary therapies has followed a trajectory that begins with a theoretical and mathematical exploration of associated eco-evolutionary models [7, 8, 9].

The clinical strategy we study here uses evolutionary rescue theory to inform the probability of tumour extinction under multiple treatment administrations or “strikes”. Although it is more usual to consider evolutionary rescue in a conservation context, the same theory is applicable when extinction is the goal, such as in bacterial infections or cancer [10]. Since an oncologist can influence the tumour environment, they can anticipate the evolutionary trajectories of cancer clones and, in theory, follow a strategy to avoid evolutionary rescue and so cure the patient [11]. The key idea is that, even if a single strike fails to eradicate cancer cells due to resistant phenotypes, it can still render the population small and fragmented. Small populations are more vulnerable to stochastic extinction and less capable of adapting to environmental changes owing to loss of phenotypic heterogeneity [10]. Cell proliferation may also slow due to Allee effects [12]. Subsequent therapeutic strikes, if well timed, can exploit these weaknesses to drive the cancer cell population to extinction [13].

Combination cancer therapies are typically designed such that cells resistant to one treatment are likely to have collateral sensitivity to another [14]. The main differences between multi-strike therapy and conventional combination therapy are in the timing of the strikes and the use of evolutionary principles to guide treatment. In combination or sequential therapy, the second or subsequent treatments are usually given during relapse, when the first treatment appears to have failed. Another conventional strategy is to simultaneously administer multiple drugs with collateral sensitivities from the beginning of treatment [15]. In multi-strike therapy, the idea is instead to attack the cancer at its weakest point when it may well be clinically undetectable. It has been suggested in a prior study that the best time to give the second strike may be while the tumour is still shrinking in response to the first therapy [16]. The success rate of multi-strike therapy is expected to be highly sensitive to the timing and severity of the second and any subsequent strikes.

Demonstrating its potential to improve cure rates across diverse cancer types, three clinical trials of multi-strike therapy are already underway. A Phase 2 trial using conventional chemotherapy drugs in metastatic rhabdomyosarcoma started recruiting patients in 2020 and is expected to run until 2026 [17]. A Phase 1 trial in metastatic prostate cancer (2022-27) involves agents that exploit the hormone sensitivity of cancer cells [18]. A Phase 2 trial using targeted therapies in metastatic breast cancer began in 2024 [19]. Further trials are in development.

Yet, despite this rapid progress to clinical evaluation, many critical questions regarding the timing of the subsequent strikes, the time until extinction, the effect of environmental and demographic factors, and most importantly, the conditions under which multi-strike therapy is a feasible alternative to other therapies remain unanswered. How effective is the first strike, and does it make the population vulnerable enough for further strikes to work? What is the probability that a population is rescued either by pre-existing mutants or those that arise during the treatment? How do outcomes vary with the cost of resistance, density dependence, and other factors that affect clonal growth rates?

We tackle these pressing questions in two ways. First, using ideas from evolutionary rescue theory, we develop and study the first analytical model of two-strike therapy. This simple, tractable mathematical model enables us to compute extinction probabilities and to identify the optimal time for the second strike. Second, we use extensive stochastic simulations to test the robustness of our analytical results and to study the effects of additional factors. We thus establish a necessary foundation for further theoretical, experimental, and clinical investigations of multi-strike therapy.

## 2 Methods

To obtain general, robust insights into the factors that determine a successful two-strike treatment strategy, we combine a deterministic analytical model and a stochastic simulation model. Both models involve two stressful environments *E*_1_ and *E*_2_ (corresponding to the two treatment strikes) and four cell types. Cells can be sensitive to both treatments (*S*), resistant to one treatment but sensitive to the other (*R*_1_ and *R*_2_), or resistant to both treatments (*R*_1,2_). The time of switching to the second treatment is *τ* and the population size at this time is *N*(*τ*).

### 2.1 Analytical methods

Our analytical modelling method is composed of two stages (Figure 1). First, we model the population dynamics during the first treatment as a set of numerically-solved differential equations. We then use those solutions to predict extinction probability using evolutionary rescue theory.

**Figure 1:**
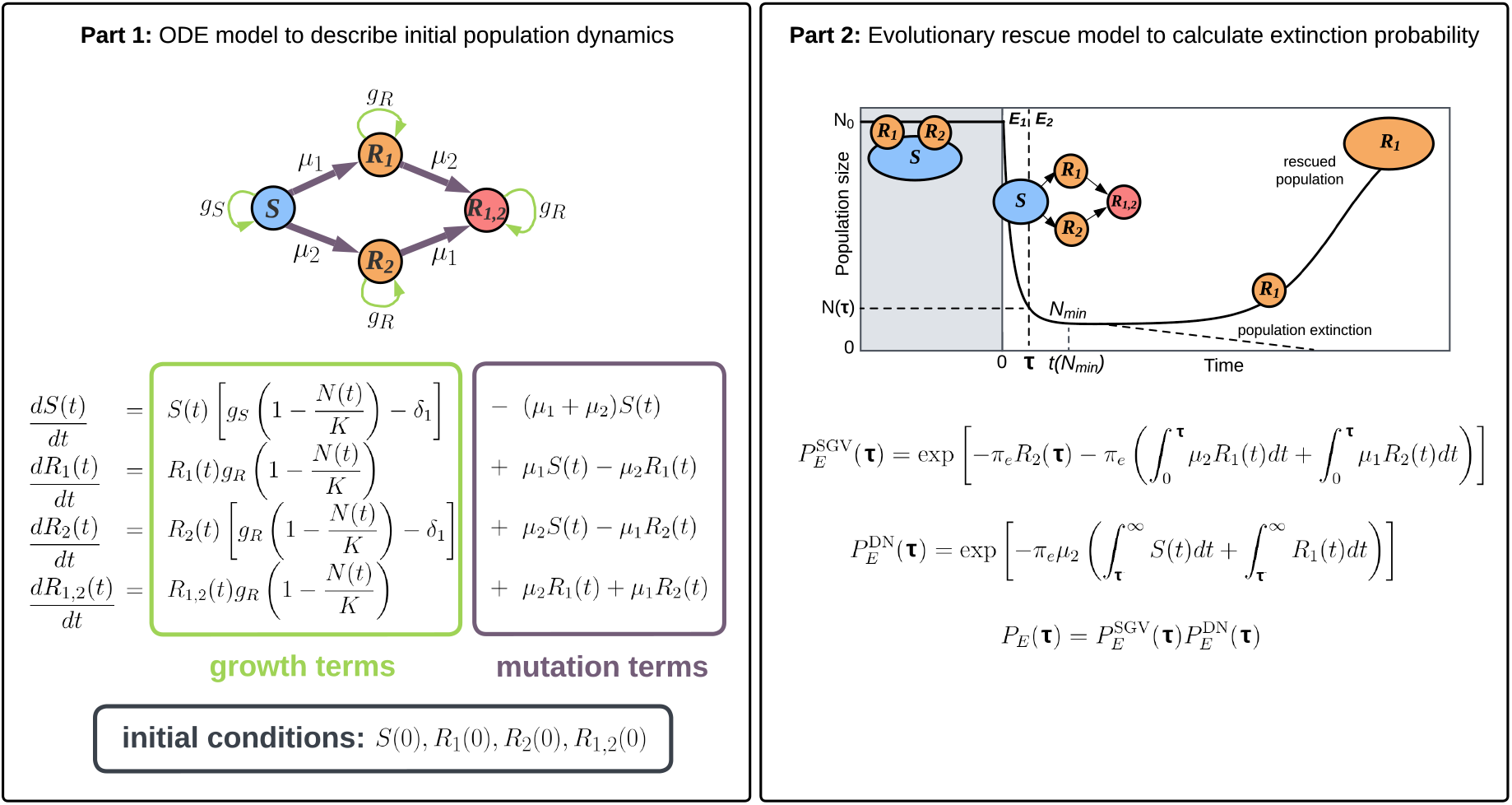
A schematic describing the deterministic analytical model. Part 1 (left) shows the ODE population growth model during the first treatment (in *E*_1_). Part 2 (right) uses input from the ODE model and evolutionary rescue theory to calculate extinction probabilities if the second treatment is given at time *τ*. The solid curve shows the population trajectory when only treatment 1 is applied. Sensitive cells are denoted by *S*. Cells resistant to treatment 1(2) and sensitive to treatment 2(1) are denoted by *R*_1_(*R*_2_). Cells resistant to both treatments are denoted by *R*_1,2_. The per capita rate of acquiring resistance to treatment 1(2) is denoted by *µ*_1_(*µ*_2_). Growth rates *g*_*S*_ for sensitive cells and *g*_*R*_ for resistant cells depend on the intrinsic birth rate, intrinsic death rate and the cost of resistance (see Table 1). The treatment-induced death rate is denoted by *δ*_1_. Initial conditions are specified by the initial population sizes of *S, R*_1_, *R*_2_ and *R*_1,2_ cells. The total initial population *N*(0) is the sum of these four subpopulations.

To calculate extinction probabilities due to standing genetic variation (pre-existing rescue mutants) and *de-novo* rescue mutants, we must obtain the population composition at the beginning of the second strike. For this we use the system of differential equations given in Figure 1 (first box), describing logistic growth in environment *E*_1_ of the four subpopulations *S*(*t*), *R*_1_(*t*), *R*_2_(*t*) and *R*_1,2_(*t*) that make up the tumour cell population *N*(*t*). The model includes mutations from less resistant to more resistant states while, for simplicity, ignoring negligible back mutations. For plausible parameter values, a tumour that grows from a single treatment-sensitive (*S*) cell is unlikely to harbour any doubly-resistant (*R*_1,2_) cells at the time it is first treated (see Appendix A.1.1). If this were not so then a cure would be highly improbable. We therefore assume *R*_1,2_(0) = 0. Other default initial conditions and parameter values are listed in Table 1. By solving the differential equations numerically over the course of the first treatment (time 0 to time *τ*), we determine the subpopulation sizes at the time of switching to the second treatment.

**Table 1:** List of parameters and initial conditions used in the analytical and stochastic simulation models, along with their default values. Note that for the analytical model, we use the values of growth rates for sensitive and resistant cells, *g*_*S*_ = *b* − *d* and *g*_*R*_ = *b* − *c* − *d*, respectively.

| Symbol | Description | Default value |
| --- | --- | --- |
| $K$ | Carrying capacity of the system | $N(0)$ |
| $b$ | Per capita birth rate of $S$ cells | 1.0 |
| $d$ | Per capita death rate of all cell types | 0.1 |
| $c$ | Cost of resistance | 0.5 |
| $\mu_1, \mu_2$ | Mutation rate for acquiring resistance to treatment 1, 2 | $2.5 \times 10^{-6}$ |
| $\delta_1, \delta_2$ | Per capita death rate due to treatment 1,2 | 2.0 |
| $S(0)$ | Initial population of $S$ cells | $10^6$ |
| $R_1(0)$ | Initial population of $R_1$ cells | 100 |
| $R_2(0)$ | Initial population of $R_2$ cells | 100 |
| $R_{1,2}(0)$ | Initial population of $R_{1,2}$ cells | 0 |

Given the population composition at treatment switching time *τ*, we next compute the probabilities 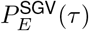 and 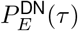 of no evolutionary rescue due to standing genetic variation and *de-novo* mutations, respectively (Part 2 in Figure 1; Appendix A.3). Because successful treatment requires the eradication of both pre-existing and *de-novo* mutants during the second treatment period, *E*_2_, the tumour extinction probability *P*_*E*_(*τ*) is the product of these two probabilities:

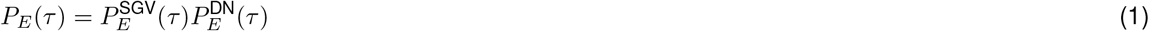

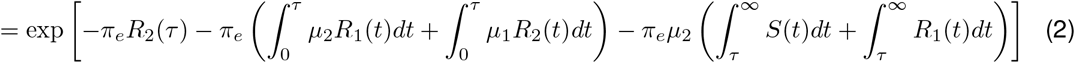

where *π*_*e*_ is the probability of establishment of a single resistant lineage, which depends on parameters *b, d* and *c*(see Appendix A.5 for the derivation). With Eq 2, we study the behaviour of extinction probability as a function of *τ* under different conditions.

### 2.2 Stochastic simulations

To test the robustness of our analytical results, we separately obtain extinction probabilities using a stochastic simulation model with the same initial conditions as the ODE system and equivalent default parameter values (see Appendix A.6). The main difference between the models is that the analytical method uses evolutionary rescue theory to calculate extinction probabilities, whereas the computational approach uses the stochastic Gillespie algorithm to simulate birth, death, and mutation events. Each simulation ends with one of three outcomes: extinction, progression, or persistence (see Table A.1). The extinction probability is estimated as the proportion of extinction outcomes in a large number of simulations.

### 2.3 Comparing results across parameter values

To compare treatment outcomes for varied parameter values, we use a summary variable *N*_*q*_ to describe how small the tumour must be at the time of switching treatment to achieve a given probability of extinction. This concept is based on our observation that the extinction probability *P*_*E*_(*τ*) generally decreases as *N*(*τ*) increases, unless *N*(*τ*) is very close to the population nadir that would pertain in the absence of a second strike (*N*_min_). Therefore, for a given extinction probability *q*(with 0 ≤*q* ≤max(*P*_*E*_(*τ*))), we can obtain a corresponding value *N*_*q*_, which is the maximum population size threshold below which we achieve an extinction probability greater than or equal to *q*. In other words, if *N*(*τ*) ≤*N*_*q*_, then we will achieve an extinction probability of at least *q*. Any given switching point *N*(*τ*) is reached twice in the trajectory of a population undergoing evolutionary rescue, once before and once after the start of relapse. *N*_*q*_ is therefore defined for both before-nadir and after-nadir switching points:

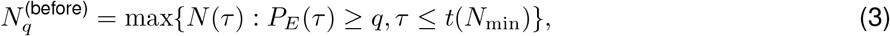

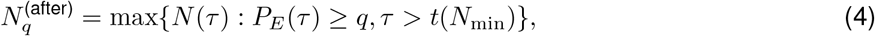

where *t*(*N*_min_) is the time at which the nadir would be reached in the absence of a second strike. *N*_*q*_ values tell us when and how fast the extinction probability drops from a high value to a low one. For instance, if the difference between 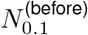 and 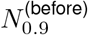 is slight, then the extinction probability must increase steeply between these two population sizes. A higher value of 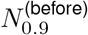 implies a wider window of opportunity for implementing a successful second strike. More generally, we want the range of *N*(*τ*) values with low *P*_*E*_(*τ*) to be as small as possible. We plot *N*_*q*_ versus *q* to analyse the trend in extinction probabilities across the range of potential switching points. Several metrics for comparing parameter sets based on *N*_*q*_ are described in Appendix A.8.

## 3 Results

To enable us to uncover general principles and determine the most important factors in a successful treatment strategy, we consider the simplest case of two strikes. Since further strikes can only increase extinction probabilities, we thus obtain conservative lower bounds on potential clinical benefits. The first treatment (or strike) creates a stressful environment that we denote *E*_1_. After switching to the second treatment, the tumour enters the second stressful environment, *E*_2_.

Corresponding to the two treatments, we consider four cell types – sensitive to both treatments (*S* cells), resistant to one of the treatments but sensitive to the other (*R*_1_ and *R*_2_) and resistant to both treatments (*R*_1,2_). Consequently, *R*_1_ and *R*_1,2_ cells are resistant in *E*_1_, and *R*_2_ and *R*_1,2_ cells are resistant in *E*_2_. Even if they initially rescue the population, all *R*_1_ cells will eventually go extinct due to the second strike. Evolutionary rescue from the second strike must be due to surviving *R*_2_ or *R*_1,2_ cells, called rescue mutants.

We obtain extinction probabilities using both a deterministic analytical model (Section 2.1) and a stochastic simulation model (Section 2.2), and we compare the two wherever possible. Unless mentioned otherwise, we use a default set of parameters and initial conditions (Table 1). Except in Section 3.4, we assume the two treatments induce identical death rates (that is, *δ*_1_=*δ*_2_=*δ*). For brevity, we will use “dose” and “treatment level” to refer to these treatment-induced death rates, which in reality also depend on pharmacodynamics and pharmacokinetics.

Our focus will be on the population size at the time of switching between the two treatments, denoted *N*(*τ*). Since the optimal *N*(*τ*) changes as we vary the parameters, the trend of extinction probabilities obtained at a fixed *N*(*τ*) could differ from the trend obtained at the optimal *N*(*τ*). The rationale for using a fixed *N*(*τ*) for such comparisons is that it may, in practice, be impossible to determine the optimal *N*(*τ*), which requires knowing the values of all the system parameters.

The fixed *N*(*τ*) can be implemented on either side of the population nadir since a given population threshold is met twice in the characteristic U-shaped trajectory of a population undergoing evolutionary rescue (Figure 2(B)). Therefore we consider both before-nadir and after-nadir switching points. The population nadir reached in the absence of a second strike, which we denote *N*_min_, can be calculated by numerically solving the system of differential equations shown in Figure 1 (see Appendix A.1.2.1 for an analytical approximation).

**Figure 2:**
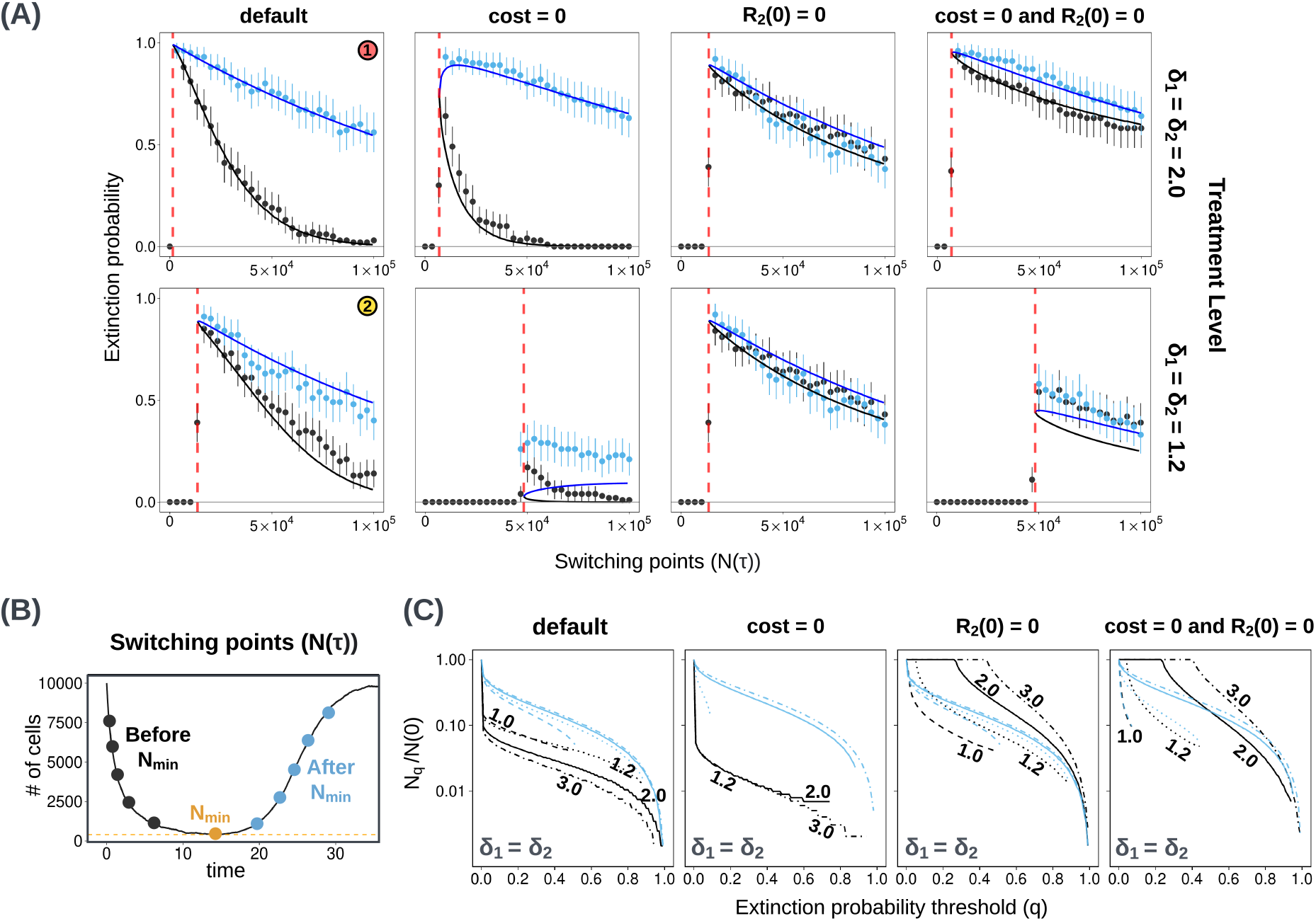
**(A)** Comparing stochastic simulation results (dots) to analytical estimates (solid line) of extinction probabilities *P*_*E*_ (*τ*) for different values of *N*(*τ*) implemented before reaching *N*_min_ (black) and after crossing *N*_min_ (blue). Red dashed lines show the expected *N*_min_ (calculated with the analytical model). Results for two treatment levels are shown (rows). Columns show results with different values for the parameter *c*(cost of resistance) and *R*_2_(0) (initial *R*_2_ population size). The default parameter set is given in Table 1. Extinction probabilities from simulations are computed using the outcomes of 100 independent runs, each with several switching points. Switching sizes smaller than *N*_min_ are equivalent to the absence of a second treatment, and result in extinction probabilities equal to zero (black points to the left of red dashed lines). Error bars show 95% binomial proportion confidence intervals. In the first column, labels (1) and (2) correspond to curves in Figure 3(C). **(B)** An illustration of switching points before and after *N*_min_, implemented with the same random seed. See Appendix A.6 for a description of the algorithm for these simulations. **(C)** *N*_*q*_ vs *q* plots for four parameter sets and different treatment levels. Black curves show before-nadir switching points, and blue curves show after-nadir switching points. Labels indicate treatment levels *δ*_1_ = *δ*_2_. The same line style (solid, dashed, dotted, mixed) is used for a given treatment level for both before and after nadir curves.

To compare different parameter values and treatment conditions, we measure the regions of high extinction probability, defined as the range of *N*(*τ*) values that give an extinction probability ≥ 0.8. Additionally, we use comparison metrics based on the quantity *N*_*q*_ (Section 2.3 and Appendix A.8), which is defined as the maximum population size threshold that must be crossed to achieve an extinction probability greater than or equal to *q*.

### 3.1 The optimal switching time is when the population size is close to its nadir

Our first aim is to find the optimal population size *N*(*τ*), at which to switch from the first to the second treatment. Our analytical and stochastic models both show that the optimal *N*(*τ*), in terms of maximizing extinction probability, is close to the population nadir *N*_min_ (Figure 2). According to the analytical model, the optimal switching point may, depending on parameter values, lie slightly before or after *N*_min_ (Appendix A.1.2.4. Yet the difference between the optimal *N*(*τ*) and the *N*_min_ is generally so small that it is not captured by our simulation results. The difference is significant only when *R*_2_ cells are initially abundant, and the cost of resistance is low (Figure 2, second column).

To explain why the optimal *N*(*τ*) is close to *N*_min_, we refer to Eq 2 and see that the maximum *P*_*E*_(*τ*) will be achieved by minimizing the sum of all the rates of generating rescue mutants. The last two integral terms are minimized slightly after *N*_min_ (see Appendix A.1.2.2 for an approximate analytical expression). However, in the terms constituting 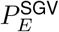 (pre-existing rescue mutants at the beginning of the second strike), the decay in the *R*_2_ population relative to the generation of *R*_1,2_ mutants determines where the optimal *N*(*τ*) lies relative to *N*_min_. For further analysis, we focus on regions of high extinction probability (*P*_*E*_ ≥ 0.8) instead of the exact optima. Consistent with our analytical predictions, we observe that these high-*P*_*E*_ regions lie around *N*_min_ (Figures A.3 and A.2).

### 3.2 It is better to implement the second strike after the nadir than before

Given the practical impossibility of treating at the exact optimal time, we next compare outcomes for treating earlier or later (see Appendix A.6 for further details of the simulation algorithm). While the optimal switching point may lie slightly before or after the nadir, we observe that switching points after *N*_min_ usually have higher extinction probabilities than those before *N*_min_ (Figure 2(A), blue versus black).

We hypothesize that this result is due the pre-existing (or initially accumulated) *R*_2_ population. If we delay switching, then this resistant subpopulation has longer to decay, which results in a smaller rescue population during the second treatment (Figure 4(A), first row, second panel). On the other hand, there is more time for doubly-resistant *R*_1,2_ mutants to accumulate. In most cases of interest, the generation of *R*_1,2_ mutants is slower than the decay of the *R*_2_ population (see Section 3.3 for exceptions), and so the window of opportunity for effective treatment extends further to the right of the nadir than to the left.

**Figure 3:**
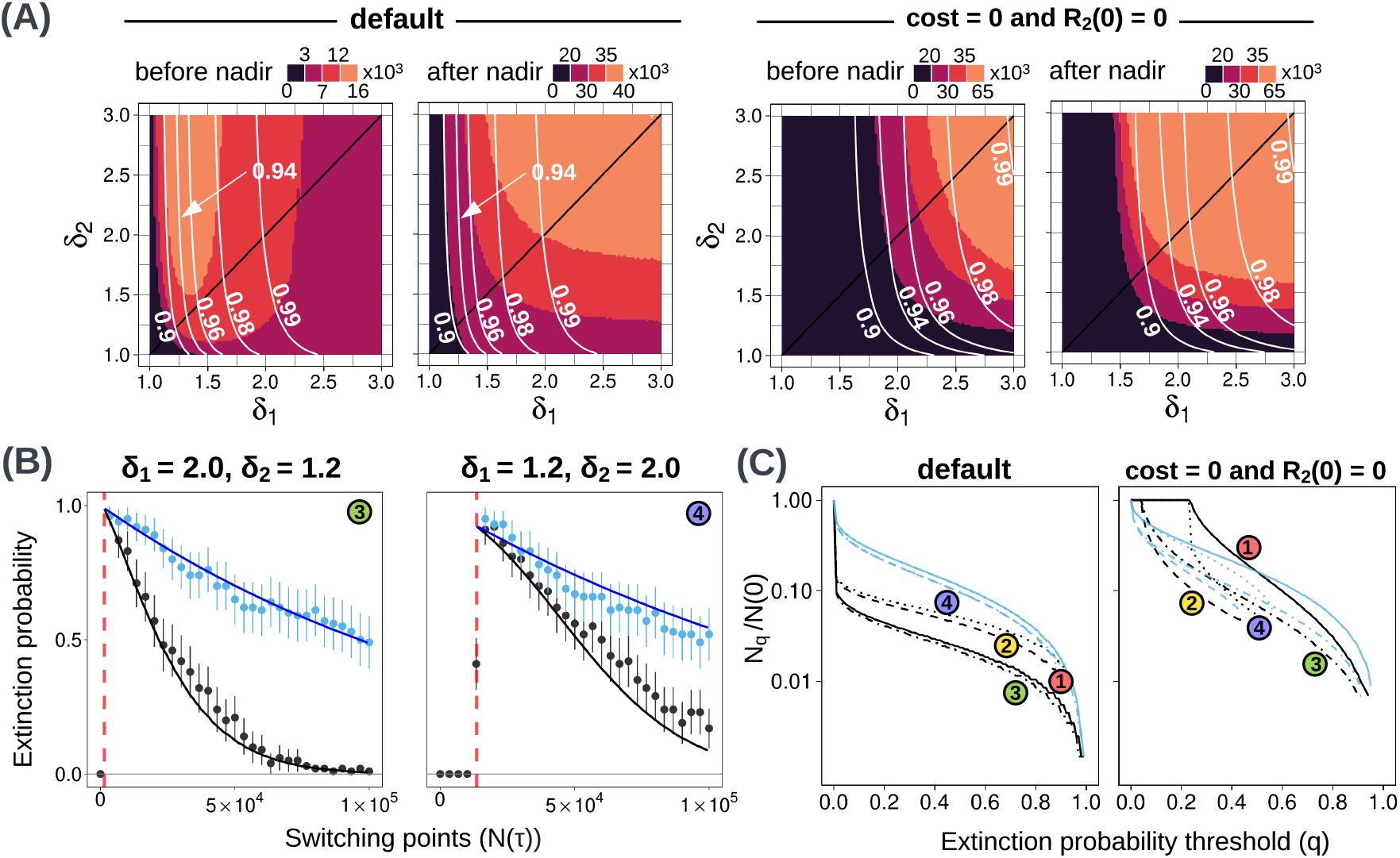
**(A)** Heatmaps (obtained from the analytical model) showing high-*P*_*E*_ regions (≥0.8) for different combinations of treatment levels *δ*_1_ and *δ*_2_. The default case is shown on the left, and the case with no cost of resistance and no initial *R*_2_ population is on the right. For each case, both before-nadir and after-nadir switching points are considered. White lines indicate optimal extinction probability contours. **(B)** Extinction probabilities for two combinations of treatment levels where *δ*_1_ ≠ *δ*_2_. Dots show simulation results and solid lines indicate analytical model predictions. Extinction probabilities are obtained from 100 paired simulations with different random seeds. Error bars show 95% binomial proportion confidence intervals. **(C)** *N*_*q*_ vs *q* plots for the default case and the case with no cost of resistance and no initial *R*_2_ population. In both cases, four treatment dose combinations are shown (labels refer to treatment combinations shown in (B) and Fig 2(A)). Black(blue) lines show before(after)-nadir switching points. The same line style (solid, dashed, dotted, mixed) is used for a given treatment level for both before and after nadir curves.

**Figure 4:**
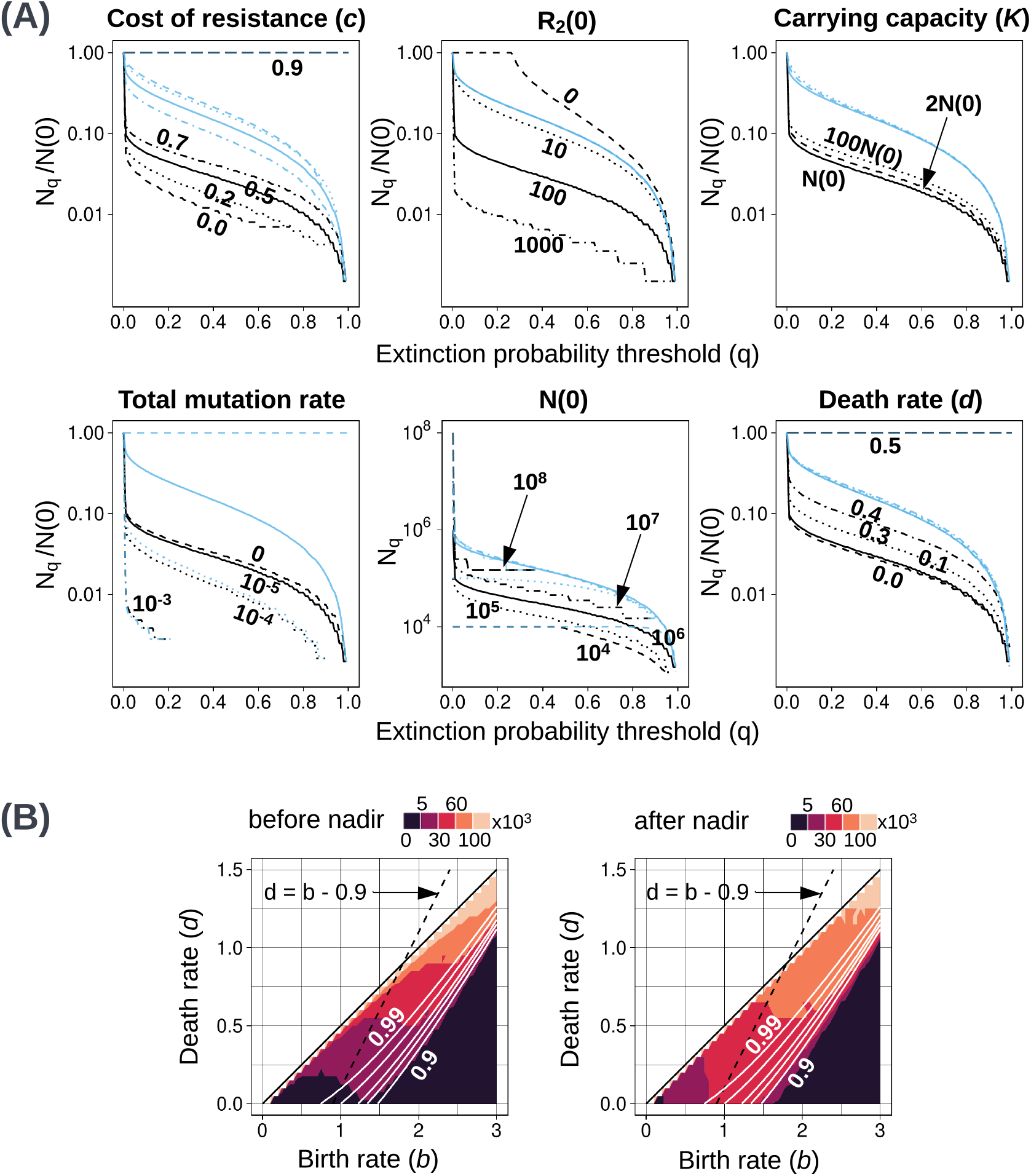
Effects of varying parameter values or initial conditions. **(A)** *N*_*q*_ versus *q* plots for several parameters and initial conditions. The *x*-axes show extinction probability threshold *q*, and *y*-axes are the (normalised) 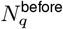 (black) and 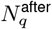 (blue) values. The only plot which is not normalised by the initial population is the one showing variation in *P*_*E*_ with *N*(0) (second row, second panel), assuming a constant proportion of initial resistant cells. The title of each plot indicates the parameter or initial condition that varies across curves. Solid curves correspond to the default parameter values (Table 1), and the same line style (solid, dashed, dotted, mixed) is used for both before and after nadir curves. In the first and last panels showing variation with changing cost of resistance and death rate, respectively, values beyond 0.9 and 0.5 are not considered since the resistant cell growth rate becomes negative. **(B)** Heatmap of high-*P*_*E*_ regions for different parameter values in *b*-*d* space. Only non-negative growth rates (excluding the effects of treatment) are considered (*d* ≤ *b* − *c*, solid black line). The dashed black line indicates the set of birth and death rates corresponding to our default growth rate (*b* − *d* = *g*_*S*_ = 0.9). Solid white lines show optimal extinction probability contours.

### 3.3 The cost of resistance and the initial *R*_2_ population size modulate extinction probabilities

Extinction probabilities depend on the number of *R*_2_ cells present at the time of switching, which in turn depends on both the cost of resistance and the number of *R*_2_ cells at the start of the first treatment. By removing one or both of the latter two factors, we can better understand how they affect treatment outcomes.

A cost of resistance is expected to hasten the decay of *R*_2_ mutations during the first treatment phase and so make the second treatment more effective. Accordingly, in most cases removing the cost of resistance reduces extinction probabilities (Figure 2(A,C), second column; Figure A.2; Figure 4(A), first row, first panel). The exception is that in the case of high-dose treatment (*δ* = 2), extinction probabilities for late switching points (well after the nadir) can be slightly higher in the absence than in the presence of a resistance cost (but the optimal extinction probability is high even for severe resistance costs). The reason for this counterintuitive result is that in the absence of a cost of resistance, the first treatment phase is shorter, which gives less time for the generation of new *R*_1,2_ mutants (see Appendix A.1.2.6 for a more detailed explanation). If we fix the switching time instead of the switching size, we see that a higher resistance cost is always beneficial (Figure 5, first and second columns). For intermediate-dose treatment (*δ* = 1.2), the main effect of removing the cost of resistance is to increase *N*_min_, which makes it impossible to achieve high rates of extinction (Figure 2(A), second row, second column).

**Figure 5:**
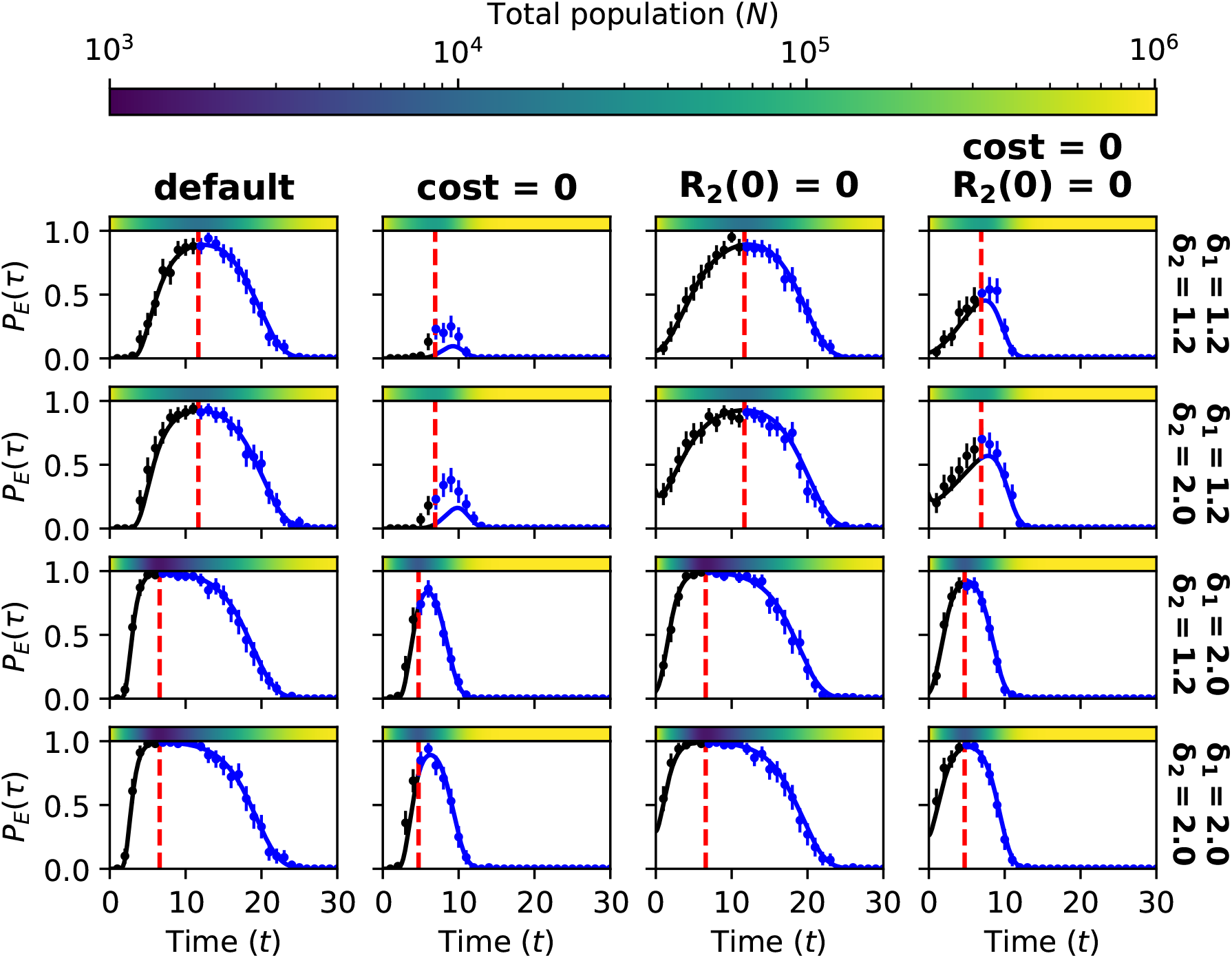
Time windows of high extinction probability (solid lines obtained from the analytical model and dots with 95% confidence intervals from 100-replicate Gillespie runs). Each subplot shows extinction probability trajectories under different modifications of the default parameter values. Population sizes are shown in the coloured bar, and the dotted red line shows when the nadir is achieved in the absence of a second strike.

Assuming that *R*_2_ cells are absent at the start of the first treatment substantially increases extinction probabilities for before-nadir switching (Figure 2(A,C), third and fourth columns; Figure A.2). The result of switching before the nadir is then similar to – or, in the case of high-dose treatment, even slightly better than – the result of switching at the same tumour size after the nadir.

Although our analytical predictions are generally very close to our simulation results, they underestimate the probability of extinction in the case of intermediate treatment dose and no cost of resistance (Figure 2(A), second row, second and fourth columns). In this case, our modeling assumption of a Poisson-distributed *R*_2_ population breaks down (see Sections A.3 and A.4 for details).

### 3.4 Higher doses do not necessarily maximize cure rates

We next relax our assumption of equal doses during the two treatment phases by examining alternative combinations of *δ*_1_ (first treatment dose) and *δ*_2_ (second treatment dose). In each case, we consider both the optimal switching point and the range of *N*(*τ*) values that lead to a high extinction probability (*P*_*E*_ ≥0.8). We will refer to the default doses 2 and 1.2 as high and intermediate, respectively.

We first consider the before-nadir regime. The intuitive prediction is that higher doses should lead to larger high-*P*_*E*_ regions. This is what we observe in the case of no resistance cost and *R*_2_(0) = 0 (Figure 3(A, C)). However, for our default parameter values, higher values of *δ*_1_ give smaller high-*P*_*E*_ regions (Figure 3(A), first row, second panel). Similarly, in the before-nadir regime, a lower dose results in higher extinction probabilities for all switching points not very close to the nadir (Figure 3(B,C) and 2(A)). A normalised *N*_*q*_ versus *q* plot for four *δ*_1_-*δ*_2_ combinations confirms that intermediate *δ*_1_ and high *δ*_2_ produce the best treatment outcome in terms of high-*P*_*E*_ regions, because it gives a higher extinction probability at the same *N*(*τ*) (Figure 3(C)). When the two doses are equal (along the black line in Figure 3(A)), we observe a similar trend (Figure A.7). Thus, for our default parameter values, an intermediate first dose paired with a high second dose gives the largest window of opportunity when switching before the nadir.

The somewhat counter-intuitive result is explained by the interaction of the dose, the cost of resistance, and the *R*_2_ population. For lower *δ*_1_, the *S* cells decay more slowly, so the switching point *N*(*τ*) is reached later. This provides more time for the *R*_2_ population to decay but also more time for *R*_1,2_mutants to arise. When the cost of resistance and the initial *R*_2_ population size are both large, the benefit of a lower *δ*_1_ outweighs the disadvantages (see Appendix A.1.2.5 for a formal explanation). We do not observe this effect when the cost of resistance or the initial *R*_2_ population size is set to zero (Figure A.4, Figure 3(A)). Note that if the dose is too low then the population size will never become small enough to permit stochastic extinction.

Although an intermediate *δ*_1_ gives a larger high-*P*_*E*_ region, the optimal extinction probability in all cases is obtained when both treatment levels are high (white contours in Figure 3(A)). Therefore, it is important to define what we need for a good treatment outcome. The best dose combination should not only lead to a high extinction probability at the optimal switching point, but it should also offer a large window of opportunity. An intermediate *δ*_1_ allows a larger high-*P*_*E*_ region than a high *δ*_1_ at the cost of compromising on the optimal extinction probability (0.99 compared to 0.94). Moreover, the absolute values of *N*(*τ*) giving a high extinction probability are also high with an intermediate *δ*_1_. This is because the *N*_min_ and the high-*P*_*E*_ region, in this case, are relatively large.

Note that this result depends on the fact that we compare high-*P*_*E*_ regions based on switching sizes. The conclusion would be different if we were thinking in terms of switching times. A larger value of *δ*_1_ is expected to lead to a larger high-*P*_*E*_ time interval (Appendix A.1.2.5). Thus, the best dose in the before-nadir regime depends on how the therapy is implemented.

Outcomes for the after-nadir regime are best when both doses are high (Figure 3(A), first row, second panel). For our default parameter values, as expected, the high-*P*_*E*_ region is also much larger for the after-nadir than for the before-nadir regime. When we eliminate the cost of resistance and the initial *R*_2_ population, the optimal treatment combinations in the before-nadir and after-nadir regimes are similar (Figure 3(A,C)).

### 3.5 Two-strike therapy is feasible only in small tumours

Using the analytical model, we compare different values of *N*_*q*_ (not normalised) for different initial population sizes *N*(0), bearing in mind that the resistant population size scales with *N*(0). We observe that the absolute values of *N*_*q*_ for *q* close to 1 do not vary by more than an order of magnitude when *N*(0) ranges over three orders of magnitude, from 10^4^ to 10^7^ cells (Figure 4(A), second row, second panel). This implies that, within this range of initial tumour sizes, a high extinction probability can be achieved by applying the second strike at a sufficiently small population size (determined by the treatment dose, growth rates, and other parameters). Nevertheless, if *N*(0) is larger than 10^8^ cells then the extinction probability never exceeds 0.4 (Figure 4(A), second row, second panel). There is, therefore, a limit on the size of tumours for which two-strike therapy is likely to succeed.

### 3.6 Mutation during treatment reduces the extinction probability

Next, we examine how ongoing mutation influences the treatment outcome. In our model, there are four types of mutation (Figure 1), and the total mutation rate is 2(*µ*_1_ + *µ*_2_). In Figure 4(A), second row, first panel, we see that increasing the total mutation rates in both *E*_1_ and *E*_2_ while keeping *N*(*τ*) and the initial frequency of resistance unchanged, results in lower extinction probability. We observe the same trend if we change the mutation rate in only one environment (Figure A.5). This effect is due to higher mutation rates resulting in a larger rescue population size and hence, a higher probability of evolutionary rescue. For a total mutation rate as high as 10^−3^, the extinction probability never exceeds 0.2. On the other hand, the benefit of decreasing the mutation rate greatly diminishes after *µ* = 10^−5^ in the before-nadir regime. In the extreme, unrealistic case of abundant pre-existing resistance and very low mutation rates, the optimal switching point would be long after *N*_min_, when the *R*_2_ population has fallen to close to zero. Therefore, there is an upper bound on the extinction probability when we switch before the *N*_min_, but extinction probabilities for the after-nadir switching points are equal to 1 (blue curve in Figure 4(A), second row, first panel).

### 3.7 Extinction probability increases with death rate and turnover

To compare treatment outcomes for different plausible combinations of birth and death rates, we plot heat maps of high-*P*_*E*_ regions in *b*-*d* space (Figure 4(B)). We observe that in the lower right region (high birth rates, low death rates), extinction probabilities are very low. This result also holds for alternative metrics for comparing parameters (Appendix A.8, Figure A.1). This leaves us with a diagonal band in the *b*-*d* space within which it is possible to attain high extinction probabilities.

Within this “good” region, we make three major observations. First, a higher death rate results in a higher extinction probability (Figure 4(A), second row, third panel). Second, as the birth rate increases, optimal extinction probability decreases and the optimal *N*(*τ*) increases (Figure A.5). However, when the birth rate (of resistant cells) is close to the death rate (*b* − *c* ≈ *d*), extinction probabilities are close to 1. The high-*P*_*E*_ regions within the “good” region remain more or less the same with changes in birth rate.

Third, we observe that the high-*P*_*E*_ regions become larger as we increase the turnover (defined as the sum *b* + *d*) while keeping the intrinsic growth rate *g*_*S*_ constant (dashed line in Figure 4(B)). Note that the cost of resistance is always a fixed fraction (0.5 by default) of the birth rate of *S* cells. It follows that when increasing turnover while keeping the growth rate *g*_*S*_ constant, the growth rate *g*_*R*_ of resistant cells decreases. This leads to a smaller rescue population, contributing to the increase in extinction probability. Another effect of turnover may relate to the establishment probability of resistant mutants. As noted in Appendix A.5, turnover appears in the expression for estimating the establishment probability *π*_*e*_. Higher turnover at a constant net growth rate *g*_*S*_ leads to a lower *π*_*e*_. If it is harder for resistant lineages to establish, then there will be fewer rescue lineages, leading to better treatment outcomes.

### 3.8 Extinction probability is insensitive to carrying capacity

As the carrying capacity is increased from *N*(0) (default value), we see a slight increase in extinction probability, but this effect saturates before *K* = 10*N*(0). This is demonstrated in Figures 4(A), first row, third panel, using the analytical model and in Figure A.7 with stochastic simulation results. Systems with a lower *K* have an extra constraint on population growth since the initial population is closer to the carrying capacity. In our model, this results in a lower decay rate for *S* cells and a higher growth rate for *R*_1_ cells. As explained in Section 3.3, the cost of resistance and the *R*_2_ population size at a given *N*(*τ*) affect the extinction probability. By default, if both factors are present, then an increase in *K* causes a slight increase in *P*_*E*_(*τ*). The individual effects of both these factors are shown in Figure A.6. The same figure shows that, in the absence of both factors, an increase in the carrying capacity results in a small decrease in *P*_*E*_(*τ*).

## 4 Discussion

We study a novel evolutionary therapy for cancer that aims to push tumours to extinction by exploiting stochasticity in small and vulnerable populations. This is done by applying multiple treatment strikes at appropriate times. A tumour that responds well to the primary therapy is primed for a second strike when it is small and susceptible to stochastic effects. The aim then is to “kick it while it’s down” [11]. This new strategy demands urgent theoretical investigation given that three clinical trials are already underway [17, 18, 19] and others are in development.

Here, we have developed the first analytical model of a two-strike therapy derived from the principles of evolutionary rescue. This model is mathematically tractable and yields clearer explanations and more general results than previous approaches in modelling “extinction therapy” [16]. We have also developed a complementary stochastic simulation model, which generally confirms the accuracy of our analytical predictions. We have sought to make both models as simple as possible, with minimal assumptions about parameter values and relations between different quantities.

We have used these new mathematical and computational models to investigate the optimal timing of the second strike and how the treatment outcome depends on crucial system parameters, including treatment levels and cost of resistance. The combination of analytical and computational analyses arms us with powerful tools to explore two-strike therapy in a wide range of scenarios, with a solid basis in eco-evolutionary theory.

### When do we get the optimal extinction probabilities?

The ability to analytically predict the optimal switching point for a large range of parameter values promises to aid the design of effective treatment schedules. By numerically solving our analytical model (Section 3.1), we have shown that the optimal *N*(*τ*) is close to *N*_min_, the population nadir in the absence of a second strike. This result – which is supported by extensive simulations (Figure A.3) – is consistent with a hypothesis proposed in the previous investigation of two-strike extinction therapy [16]. In contrast to the previous study, which suggested that striking before the *N*_min_ is better, our results showed the optimal switching point may be slightly before or after the nadir (Section 3.1 and Appendix A.1.2.4).

Since it is unreasonable to expect switching to the second strike exactly at the optimal point, we examined high-*P*_*E*_ (≥0.8) regions or windows of opportunity. We found that switching after the nadir generally results in a higher extinction rate than switching before (Section 3.2) because the window of opportunity extends further into the after-nadir regime. The difference in the extinction probabilities between before and after nadir switching points depends on the cost of resistance and the size of the initial population resistant to treatment 2 (Section 3.3). We conclude that it is generally better to wait slightly longer and risk missing the optimal *N*(*τ*) than to apply the second strike too early. However, one should certainly not wait until the tumour becomes detectable again (as is the current practice) because that increases the probability that rescue mutants will emerge.

In practice, the tumour detection thresholds are much higher than the initial population sizes we consider in our model (∼1 million cells, see Section 3.5). Therefore, it is important to have an estimate of the population size as well as time windows in which switching to the second treatment gives a high extinction probability. In Figure 5, we compare the high-*P*_*E*_ time window to the total duration in which the tumour is undetectable and see that the time window sizes correlate with the size of high-*P*_*E*_ regions in most cases. A notable exception is when the cost of resistance is zero and the treatment level is high. In this case, we observed a larger high-*P*_*E*_ region but a shorter high-*P*_*E*_ time window compared to the case with a high cost of resistance (Figure 2(A), first row and Figure 5, fourth row). For clinical applications, the size of the window of opportunity, both in terms of time and population size, should be considered.

### What are the optimal doses?

The treatment levels during the two strikes (*δ*_1_ and *δ*_2_) are the easiest model parameters to control in practice. The higher the two doses, the higher the extinction probability at the optimal switching point (Figure 3.4). However, switching at the optimal size, which becomes smaller as the treatment dose is increased, may not be feasible. In this case, the treatment combination that gives a wide range of switching points with a high probability of extinction may be better. Surprisingly, at least with a high cost of resistance, we found that the largest high-*P*_*E*_ (≥ 0.8) region in the before-nadir regime is obtained with an intermediate first dose paired with a high second dose (Section 3.4).

This result emphasizes the importance of timing in two-strike therapy – a stronger treatment with a poorly chosen switching time can be worse than a weaker treatment given at the right time. An interesting implication of this result is that the two treatments need not both be very effective. An intermediate treatment-induced cell death rate can give good treatment outcomes if the cost of resistance is high. Moreover, the optimal switching threshold is also relatively high for a less effective first treatment, which may be beneficial in practice.

### What other tumour parameters determine the success of two-strike therapy?

Our systematic exploration of the model parameter space reveals several noteworthy effects on treatment outcomes. First, although a high cost of resistance is predictably beneficial, we found that two-strike therapy can outperform conventional treatment even when this cost is small or non-existent (Section 3.3). Therefore, in common with adaptive therapy [20], two-strike therapy is not contingent on a cost of resistance. Moreover, we saw a variable response to a change in the cost of resistance depending on treatment level. For high treatment doses, we observed comparable extinction probabilities in the presence and absence of the cost of resistance, but for intermediate doses, a small cost of resistance gives much worse treatment outcomes than a high cost.

Second, mutation during either treatment is detrimental to treatment outcome (Section 3.6). This result suggests, for example, that mutagenic therapies may be less appropriate.

Third, we find that higher death rates and higher turnover are beneficial, as has previously been shown for adaptive therapy [21]. Fourth, although a higher carrying capacity allows more tumour growth, we found that for a given initial tumour size, changes in carrying capacities have little effect on treatment outcome (Section 3.8).

Although we have used simple models with minimal assumptions to ensure that our main findings are qualitatively robust, we have not explored all plausible functional forms. For example, the effect of changing the mutation rate might be different in a model in which mutations occur only at the time of cell division. Therefore, our results are subject to certain methodological assumptions and limitations.

### When should two-strike therapy be used?

Two-strike therapy holds most promise as an alternative to conventional therapy in cases where a very good initial response to treatment is typically followed by relapse. Our results suggest it is likely to succeed only in relatively small tumours (Section 3.5). Nevertheless, we expect that subsequent treatment strikes, following the same principle, would lead to higher extinction probabilities for larger tumours. Additionally, we have assumed a larger initial resistant population than expected for our default tumour size (see Appendix A.1.1), providing conservative estimates of extinction probabilities. Including an Allee effect [12] is also expected to increase extinction probabilities, making the therapy viable in a wider range of scenarios [16]. Hence even two-strike therapy may be feasible in substantially larger tumours.

Two-strike therapy may be a wise strategy when one of two available treatments is less effective than the other. Conversely, if resistant cells are abundant and have relatively high fitness, then this therapy is unlikely to succeed, and a long-term tumour control strategy such as adaptive therapy could be a better option [9, 20]. Even when it may be theoretically optimal, two-strike therapy crucially depends on the availability of effective treatments with low cross-resistance and methods for monitoring tumour burden over time [22].

Demonstrating the broad feasibility of two-strike therapy, the three clinical trials that are already underway in metastatic rhabdomyosarcoma [17], metastatic prostate cancer [18], and metastatic breast cancer [19] involve not only diverse cancer types but also very different classes of treatment, including chemotherapy, targeted therapies, and hormonal agents. Other proposed targets include locally advanced rectal adenocarcinoma [23] and paediatric sarcomas [22].

## Conclusion and future directions

We have shown that two-strike therapy is a theoretically sound concept that, in certain scenarios, could plausibly increase cancer cure rates. Our work provides a necessary foundation for further mathematical investigation and justification for experimental and clinical testing of this innovative strategy.

An important topic for further mathematical analysis is the prevention of evolutionary rescue with more than two strikes. Previous work on the optimal scheduling of multiple treatments [24, 25, 26] suggests that alternating two treatments is a theoretically sound approach. An alternative strategy, more in line with the original conception of extinction therapy, is to switch to a third treatment whenever possible. Other immediate directions for mathematical investigation include accounting for cross-resistance and considering alternative biological assumptions, such as modelling resistance as a continuous, plastic trait.

## Code availability

All relevant code is available in a public repository [27].

## Funding and acknowledgements

SP and RN were supported by the National Cancer Institute of the National Institutes of Health under Award Number U54CA217376. YN benefited from the European Union’s Horizon 2020 research and innovation programme under the Marie Skłodowska-Curie grant agreement No 955708. YN and RN were also supported by a London Mathematical Society Research in Pairs award (reference 42320). The opinions expressed in this document reflect only the author’s view and in no way reflect the European Commission’s opinions. The European Commission is not responsible for any use that may be made of the information it contains. The content is solely the responsibility of the authors and does not necessarily represent the official views of the National Institutes of Health. All simulations were performed on City, University of London’s Hyperion cluster.

## Contributions

RN conceived the research question. RN and YV supervised the project. SP and RN designed the research. SP developed the models. SP and AA ran simulations. SP, AA and YV carried out the mathematical analysis. SP wrote the paper with contributions from AA, YV and RN. All authors approved the manuscript.

## A Appendices

### A.1 Analytic Model Without Competition

We study here the model without competition, obtained by letting the carrying capacity *K* go to infinity. This is a reasonable approximation of the true model (Figures 4A and A.7, bottom right subplot), and it may be solved explicitly. This allows us to better understand the impact of various parameters.

We denote by *g*_1_, *g*_2_, and *g*_1,2_ the baseline per-cell growth-rates of the resistant strains *R*_1_, *R*_2_ and *R*_1,2_, respectively. In the main text, for simplicity, these growth rates are assumed to be equal and denoted by *g*_*R*_, but this assumption is not needed here. Let *γ*_*S*_ = *g*_*S*_ − *µ*_1_ − *µ*_2_, *γ*_1_ = *g*_1_ − *µ*_2_ and *γ*_2_ = *g*_2_ − *µ*_1_ denote the per-cell growth-rates of subpopulations *S, R*_1_, and *R*_2_, net of outgoing per-cell mutations rates. Let *c*_*i*_ = *γ*_*S*_ − *γ*_*i*_ (*i* = 1, 2) denote the cost of resistance net of the mutation rates. Since the mutation rates are assumed to be much lower than the growth rates, *γ*_*S*_ ≃ *g*_*S*_, *γ*_*i*_ ≃ *g*_*i*_, and *c*_*i*_ ≃ *g*_*S*_ − *g*_*i*_, *i* = 1, 2.

During the administration of treatment 1, the model reads:

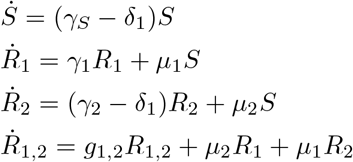

This system of linear differential equations is easy to solve. Assuming *γ*_2_ ≠ *γ*_*S*_, we get:^1^

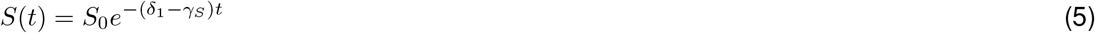

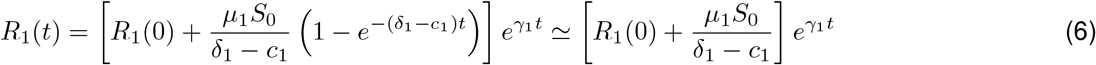

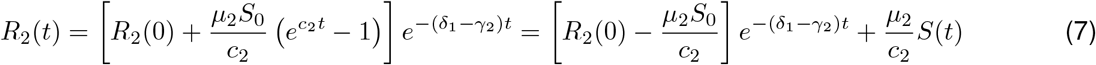

where the approximation for *R*_1_ holds for *t* large enough. The quantity *R*_1,2_(*t*) is also easy to compute, but is not relevant. Indeed, it represents the expected number of *R*_1,2_ cells, while we are interested in the expected number of potential *R*_1,2_ rescue *mutants*, i.e, of *R*_1,2_ cells directly arising from mutations from *R*_1_ or *R*_2_ cells, as opposed to reproduction of previous *R*_1,2_ cells. The latter is given by

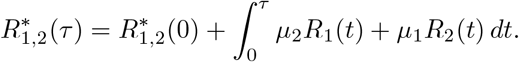

Here,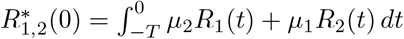 where −*T* is the time of tumour initiation.

Explicitly,

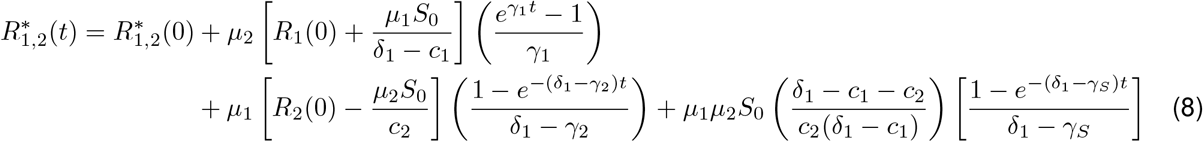

where the most important terms are those of the first line.

Letting *π*_1_ and *π*_1,2_ denote the probability of establishment of *R*_2_ and *R*_1,2_ mutants, respectively, the total expected number of rescue mutants is given by

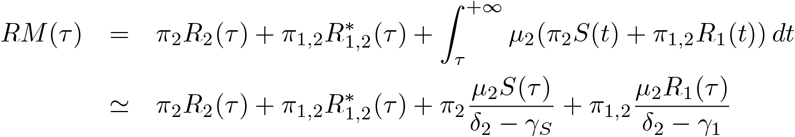

where the approximation neglects the impact of mutations from *S* to *R*_1_ on the dynamic of *R*_1_ cells during the administration of treatment 2. The expected number of rescue mutants is the quantity that we seek to minimize (assuming that, in the stochastic analogue of the model, the number of *R*_2_ cells at time *τ* is approximately Poisson distributed, see Appendix A.4).

Letting 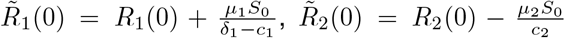, and using (7) and (8), we obtain the following approximation of the expected number of rescue mutants when 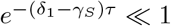:

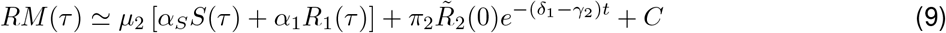

where

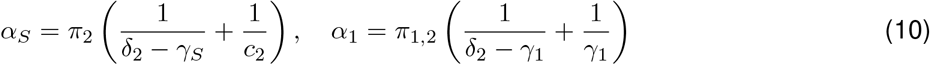

and

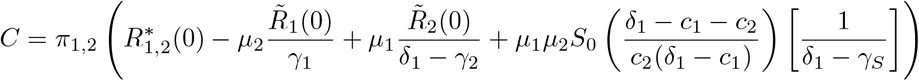

is a constant that impacts the probability of extinction but not the comparison between switching times.^2^

#### A.1.1 A reference case for initial resistant populations

The initial resistant populations depend on the tumour and treatment history before the phase that we are interested in. For instance, if other treatments were given before and that resistance to these treatments is correlated to resistance to treatments 1 and 2, the initial resistant populations *R*_1_(0) and *R*_2_(0) could be large. Nevertheless, as a reference point, assume that the tumour starts at time −*T* with a single sensitive cell and no resistant cells, and that, until the administration of treatment 1, the tumour follows the dynamics we assumed, without treatment kill rate: 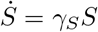 and 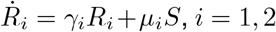. Using that 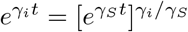, it is then easy to see that

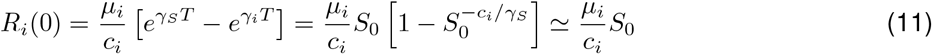

This is equal to 5 with our default parameter values, much lower than the 100 initial *R*_*i*_ cells we assume.^3,4^ Integrating the analog of (11) between −*T* and 0 leads to:

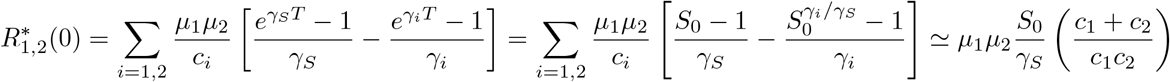

Plugging the initial resistant populations obtained in (11) in the expressions for *R*_*i*_(*t*) and 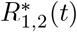 leads to (during treatment 1):

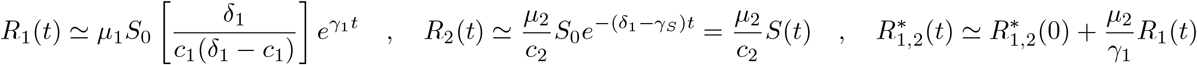

This leads to:

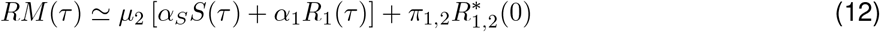

where *α*_*S*_ and *α*_1_ were defined in (10). The key difference with (9) is that, since 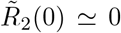, the term in 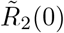 disappeared, and the *R*_2_ population is fully hidden in a part of the term *α*_*S*_*S*(*τ*). This allows us to study the worst-case scenario in which the treatment history leads to a large resistant population, giving us conservative bounds on the extinction probability.

#### A.1.2 Intuition about our findings, robustness

We now use previous formulas, with or without assuming (11), to compute some interesting quantities and get some intuition about our findings in the main text.

We first need a lemma.

##### Lemma A.1.

*Let A, B, α, β be positive real numbers. The quantity x*(*t*) = *Ae*^−*αt*^ + *Be*^*βt*^ *is minimal at* 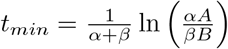 *and its value is then*

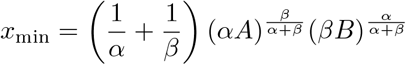

*Moreover, if C >* 0, *the function f*(*t*) = *Be*^*βt*^*/Ae*^−*αt*^ *satisfies f*(*t*) = *Cf*(*t*_*min*_) *when* 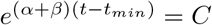, *and we then have*

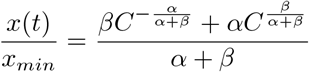

*Proof*. Let *y*(*t*) = *Ae*^−*αt*^, *z*(*t*) = *Be*^*βt*^ so that *x*(*t*) = *y*(*t*) + *z*(*t*). The function *x* is convex and is thus minimal when its derivative cancels. Since *x*^*′*^(*t*) = −*αy*(*t*) + *βz*(*t*), this occurs when *z/y* = *α/β*. This leads to 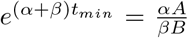, hence the formula for *t*_*min*_. Moreover, *z/y* = *α/β* implies *y/x* = *β/*(*α* + *β*) and *z/x* = *α/*(*α* + *β*). Since 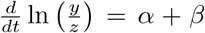, we get *f*(*t*) = *f*(*t*_*min*_)*e*^(*α*+*β*)Δ*t*^ where Δ*t* = *t* − *t*_*min*_. Thus, *f*(*t*) = *Cf*(*t*_*min*_) when *C* = *e*^(*α*+*β*)Δ*t*^. We then have 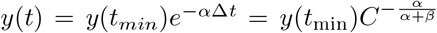 and similarly, 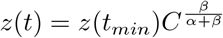. Therefore,

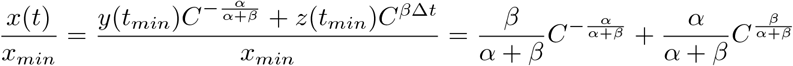

□

##### A.1.2.1 Computation of the nadir

Around the nadir, most tumour cells are either *S* cells or *R*_1_ cells,

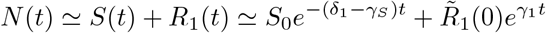

The right-hand side may be minimized using Lemma A.1. Letting 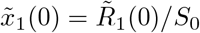, this leads to:

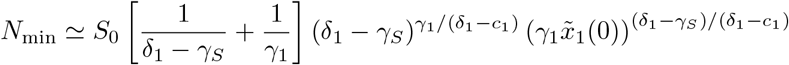

where we used that *δ*_1_ − *γ*_*S*_ + *γ*_1_ = *δ*_1_ − *c*_1_. With our default parameter values, we obtain *N*_*min*_ ≃ 2018. With the initial *R*_1_ population given by (11) and our default parameter values otherwise, we would get *N*_*min*_ ≃286. Adding competition, as in the main text, would somewhat decrease these numbers since this reduces all net growth rates.

##### A.1.2.2 The expected number of de novo rescue mutants is minimized by switching slightly after the nadir

This part of the analysis is also valid for the model with competition, as long as *N*_*min*_ ≪ *K*. Let *α* = *δ*_1_ − *γ*_*S*_ and *β* = *γ*_1_. At relevant switching times, 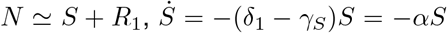 and 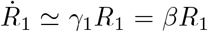, so 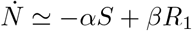. The nadir is thus reached at the time *t*_min_ such that *R*_1_*/S* = *α/β*.

Let *a*_1_ = *π*_1,2_*/*(*δ*_2_ − *γ*_1_) and *a*_*s*_ = *π*_2_*/*(*δ*_2_ − *γ*_*S*_). The expected number *λ*^*DN*^ of de novo rescue mutants satisfies

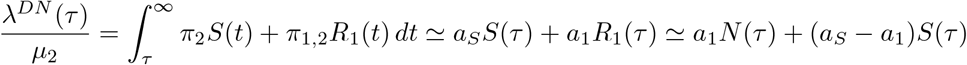

where we used that close to the nadir, *N* ≃ *S* + *R*_1_. In the absence of costs of resistance, that is, if *γ*_1_ = *γ*_2_ = *γ*_1,2_ = *γ*_*S*_ (which also implies *π*_2_ ≃*π*_1,2_ and hence *a*_*S*_ = *a*_1_), the quantity *λ*^*DN*^ (*τ*) is minimized at the nadir. If, however, we assume a cost of resistance for *R*_1_ cells and similar probabilities of establishments for *R*_2_ and *R*_1,2_ cells (*π*_2_ ≃ *π*_1,2_), then *a*_*S*_ *> a*_1_. Since at the nadir, 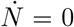 and 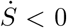, it follows that *λ*^*DN*^ is still decreasing. To minimize it, it is thus best to switch after the nadir.

To be more precise, note that *λ*^*DN*^ is minimal at the time *t*_*DN*_ such that 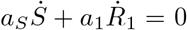, that is, −*a*_*S*_*αS* + *a*_1_*βR*_1_ = 0.. It follows that:

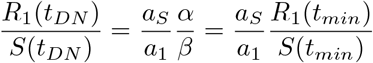

Using Lemma A.1 with 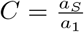 thus implies that

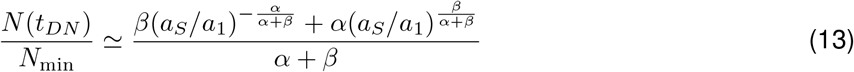

With our standard parameter values, we get *N*(*t*_*DN*_) ≃1.013 *N*_min_. The expected number of de novo rescue mutants is minimized by switching just slightly after reaching the nadir. Testing other parameter values suggests that this conclusion is robust, in the sense that the ratio (13) varies relatively little from 1 when changing parameters (Figure A.8).

##### A.1.2.3 The expected number of pre-existing rescue mutants may be minimized before or after the nadir

During treatment 1, the *R*_2_ population decreases, but the expected number of *R*_1,2_ rescue mutants increases. The expected number 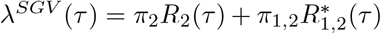 of pre-existing rescue mutants is minimal when these two forces exactly balance each other, i.e. when:

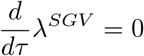

that is,

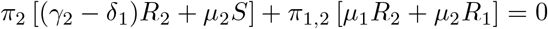

Assuming *µ*_1_ ≪ *δ*_1_ − *γ*_2_, this happens when:

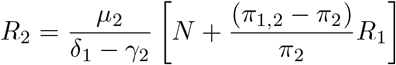

If, as in the main text, *γ*_1,2_ = *γ*_2_, hence *π*_2_ = *π*_1,2_, this boils down to:

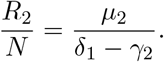

This may occur before or after the nadir, depending on the value of parameters, and in particular of the initial *R*_2_ population: the larger *R*_2_(0), the more important it is to wait that the *R*_2_ population decreases.

In the reference case 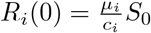, the expected number of preexisting rescue mutants is approximately

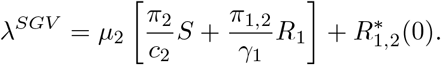

Assuming 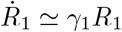 this is minimal when

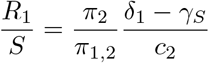

Recalling that at the nadir, 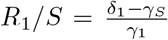, and that the ratio *R*_1_*/S* increases with time, it follows that the number of preexisting rescue mutant reaches its minimum before the nadir if

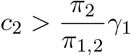

and after the nadir otherwise. With our default parameters, the cost of resistance is very strong, and this inequality is satisfied (0.5 *>* 0.4). Thus, the number of preexisting rescue mutants reaches its minimum before the nadir. More precisely, by Lemma A.1 and using 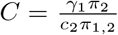, the expected number of pre-existing rescue mutants is minimized at the time *t*_*SGV*_ such that *t*_*SGV*_ ≃ 0.983 *t*_*min*_, and the tumour size is then *N*(*t*_*SGV*_) ≃ 1.005*N*_*min*_.

However, with a smaller cost of resistance *c*_2_, the number of preexisting rescue mutants would reach its minimum after the nadir. The optimal switching time would then unambiguously be after the nadir, even though we assumed in this computation a much smaller initial *R*_2_ population than in our simulations. Thus, the fact that the optimal switching time is after the nadir does not require a large initial *R*_2_ population.

##### A.1.2.4 When to switch in total: before or after the nadir?

To see whether the optimal switching time is before or after the nadir, consider first the reference case where 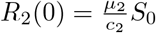. Since at relevant switching times, *N* ≃ *S* + *R*_1_, Eq. (12) may be rewritten:

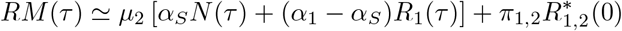

Since at the nadir *dN/dτ* = 0, and *R* is increasing, it follows that *dRM/dτ* has then the sign of *α*_1_ − *α*_*S*_. Thus, it is best to switch after the nadir if *α*_1_ *< α*_*S*_, that is, if:

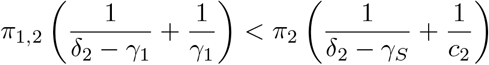

This condition may be understood as follows: to minimize the number of de novo rescue mutants, it is best to switch after the nadir if 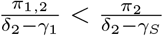. To minimize the number of pre-existing rescue mutants, it is best to switch after the nadir if 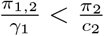. If both conditions hold, or if they hold in an appropriate average sense, then it is globally best to switch after the nadir.

With our default parameter values except for the initial resistant populations, *α*_*S*_*/α*_1_ ≃ 0.93 < 1, so it is best to switch a bit before the nadir. More precisely, from (12), we get that the expected number of rescue mutants is minimized when 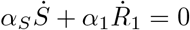, which leads to

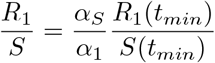

Denoting by *t*_*opt*_ the time at which the expected number of rescue mutants is minimized, we then get from Lemma A.1 with 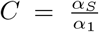 that *t*_*opt*_ ≃ 0.99994 *t*_*min*_ and that *N*(*t*_*opt*_) ≃ 1.0005 *N*_*min*_. Thus, the optimal switching time is very slightly before the nadir, but so slightly that this could be due to approximation errors in our formulas, and could be easily reversed by slight changes in parameters.^5^

We conclude that whether the optimal switching time is before or after the nadir is not a robust finding, and depends on parameters. What seems robust, however, is that it is best to switch close to the nadir. Since the time at which the nadir is reached may not be observed in practice, it is important to note that due to the assumed cost of resistance, tumour size and the number of expected rescue mutants tends to decrease relatively quickly before the nadir, and then to rebound more slowly.^6^ This makes switching after the nadir a safer move (see Figure 5).

For arbitrary initial resistant populations, replacing (12) by (9) leads to:

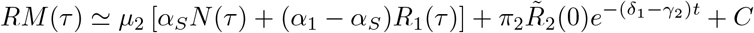

The previous argument shows that, with our default parameter values, the bracket is minimized very slightly before the nadir. However, in our simulations, we chose a relatively large initial *R*_2_ population, hence 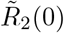 is quite positive. Since the term 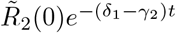 decreases with time, switching after the nadir should give better results than switching before. This is indeed what we observed.

##### A.1.2.5 A lower value of *δ*_1_ may reduce the expected number of rescue mutants when switching before the nadir and at a given size

For a given switching time, and not taking into account treatment toxicity, a larger value of treatment 1 kill-rate *δ*_1_ is unambiguously beneficial: i t leads to a lower sensitive population, hence also to fewer mutations from sensitive to resistant cells, and lower resistant populations.It thus leads to a lower probability of evolutionary rescue.^7^

For a given switching size, the situation is different, at least when we switch before the nadir, and the initial *R*_2_ population is relatively large. In that case, as mentioned in the main text, lower values of *δ*_1_ provide an advantage by letting the *R*_2_ population decay more between the beginning of treatment 1 and the time when the switching tumour size is reached. We now explain this formally.

Assume that the protocol is to switch treatment the first time that tumourreaches size 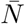, and consider various values of *δ*_1_ such that this happens. As explained above, a lower value of *δ*_1_ leads to a lower rate of decay of the sensitive population, hence also to more mutations from *S* to *R*_*i*_ and a higher growth rate of the resistant populations. It thus leads to a larger resistant population size and a larger tumour size for a given sensitive population size. It follows that to reduce tumour size to size 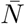, the sensitive population must be reduced by a larger factor when *δ*_1_ is low. Thus, letting *τ*(*δ*_1_) denote the time at which the switching size 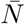 is reached, the ratio *S*(*τ*(*δ*_1_))*/S*_0_ is increasing with *δ*_1_ (lower for a low value of *δ*_1_).

But 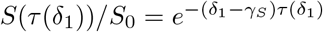, so

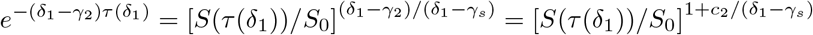

The lower *δ*_1_, the lower the quantity in the bracket, and the larger the exponent (assuming a cost of resistance). Thus a low value of *δ*_1_ leads to a low value of 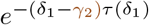, hence to a low value of the term 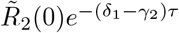 when 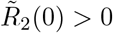.

Since a low value of *δ*_1_ increases the proportion of *R*_1_ cells when reaching size 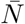, the value of *δ*_1_ also affects the term *α*_*S*_*S* + *α*_1_*R*_1_ = *N* [*α*_*S*_ + (*α*_1_ − *α*_*S*_)*R*_1_*/N*]. However, as long as the ratio *α*_*S*_*/α*_1_ is close to 1, as with our default parameter values, this effect will be small.

##### A.1.2.6 Impact of a cost of resistance

A key benefit of costs of r esistance, i f they are incurred even before treatment b egins, i s to decrease the initial proportion of resistant cells. This was addressed when deriving (11). By contrast, we focus here on the impact of costs of resistance for fixed initial resistant populations.

For a given switching time, all costs of resistance are beneficial. They do not impact the sensitive population, and reduce resistant populations, hence the expected number of rescue mutants. It follows that any cost of resistance increases the range of switching times leading to a large probability of extinction.

For a given switching size, resistance costs for *R*_2_ and *R*_1,2_ mutants are still beneficial: they do not significantly affect the switching time, nor the proportion of *R*_1_ and *S* cells at that time, but reduce the expected number of rescue mutants, through a quicker decay of the *R*_2_ population in the first phase, and the reduction of the probability of establishment of potential rescue mutants in both phases.

The effect of a cost of resistance for *R*_1_ cells when switching at a given size is more subtle. The coefficient *α*_1_ in (9) is equal to:

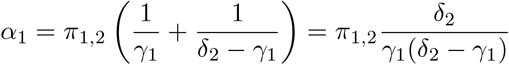

In our model, the number of mutations from *R*_1_ to *R*_1,2_ cells is proportional to the *R*_1_ population but not to its net growth rate. As a result, for a given resistant population *R*_1_(*τ*), *R*_1_ cells will generate more *R*_1,2_ mutants in phase 1 when incurring a large cost of resistance.^8^ This is reflected in the fact that the term 1*/γ*_1_ is a decreasing function of *γ*_1_, hence an increasing function of *c*_1_. They will, however, generate fewer *R*_1,2_ mutants in phase 2, as a higher cost of resistance means a quicker decay of the *R*_1_ population. This is reflected in the fact that the term 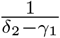 increases with *γ*_1_. The net effect depends on whether *γ*_1_ is smaller or larger than *δ*_2_*/*2 (indeed, the denominator *γ*_1_(*δ*_2_ − *γ*_1_) is maximal when *γ*_1_ = *δ*_2_*/*2). With our default parameter values, *γ*_1_ *< δ*_2_*/*2, and so the dominant effect is that a larger cost of resistance *c*_1_, hence a lower growth-rate *γ*_1_, leads to more *R*_1,2_ mutants in phase 1.

When switching long after the nadir, the increase of *α*_1_ is the main effect of a cost of resistance *c*_1_. Indeed, pre-existing *R*_2_ mutants should be rare, and the fraction of *R*_1_ cells when switching is anyway close to 1, hence should vary little with *c*_1_. A larger cost of resistance is then paradoxically detrimental.

When switching before the nadir, other effects should be taken into account: a larger cost of resistance means that it takes a bit more time to reach the switching size, giving a bit more time for the *R*_2_ population to decay (and in particular decreasing the term in 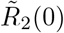 in (9) when 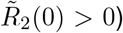. This effect will be smaller long before the nadir, as the time at which the switching size is reached then depends little on the *R*_1_ population. A larger cost of resistance also leads to a lower proportion of resistant cells when switching, which affects the term *α*_*S*_*S* + *α*_1_*R*_1_, though, as said before, this effect should remain small when the ratio *α*_*S*_*/α*_1_ is close to 1. We conclude that, though the net effect depends on parameter values, a larger cost of resistance *c*_1_ might counter-intuitively decrease the probability of survival when switching at a given size, also when switching before the nadir.

### A.2 Using results from evolutionary rescue theory

While the conventional goal of evolutionary rescue is to maximize the probability of rescue, the same mathematical formulations can be applied with the aim of extinction. For our deterministic analytical model, we use the theory of evolutionary rescue to find the probability of extinction in cancer populations undergoing two-strike therapy. A population is rescued from extinction when one or more resistant lineages are established in the population.

We assume that the generation of resistant mutants in the population can be approximated by a Poisson process [10] (see Appendix A.4 for further investigation into the validity of this assumption in the context of our model). The resistant mutants that lead to evolutionary rescue are called rescue mutants. The probability of establishment of a resistant mutant can be approximated by Haldane’s 2*s* [28], where *s* is the selective advantage of the mutant over the wild-type, which depends on the degree of stress. For a more general, continuous-time estimate, we use diffusion approximation to get the establishment probabilities of resistant lineages starting with a single cell (derived in Appendix A.5). The establishment probability is multiplied by the rate of generation of resistant mutants to obtain the rate of generation of rescue mutants. Therefore, once the rate of the Poisson process is calculated, it is easy to obtain the probability of extinction, which will be equal to the probability that no rescue mutants are generated.

We assume density independence of resistant lineages and use branching processes to model the emergence of new resistant (and rescue) mutants [29]. The growth model we use is described in Appendix A.5. Density independence is a reasonable assumption because rescue lineages become important only when the population is much smaller than the carrying capacity. Additionally, we make the assumption that the probability of establishment of a resistant mutant is constant throughout. There is extensive literature on what happens when these assumptions do not hold. Analytical results can be derived in all such situations using stochastic methods and taking continuous time approximations [30, 31, 32].

An important distinction is that the probability of rescue by pre-existing mutants (from before the onset of stress, called standing genetic variation) is different from that of new mutants (via de-novo mutations) [29]. This is because the pre-existing mutant (SGV) lineages typically get more time to grow than the de-novo mutants (DN). There are several expressions by different authors [31, 30, 29] for the probability of evolutionary rescue following a single change in environment (treatment strike in our case) by both these classes of mutants, but considering the common conceptual basis, they all reduce to the following form:

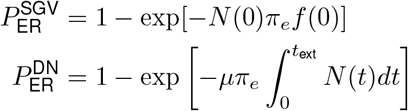

where *N*(0) is the population size at the onset of stress (*t* = 0), *µ* is the total per capita, per unit time mutation rate after the onset of stress, *f*(0) is the frequency of resistant variants at *t* = 0 and *t*_ext_ is the time at which the population would go extinct if it is not rescued. Thus, the first and second terms, respectively, represent evolutionary rescue by a population that only becomes resistant to the stress induced at *t* = 0 and evolutionary rescue by a rescue population generated by mutation from *N*.

In Section 2.1, we use these results extensively in the context of two-strike therapy. The principles of evolutionary rescue provide a foundation for building the theoretical formulation of two-strike therapy. There are, of course, significant differences to account for — firstly, most evolutionary rescue models consider either one abrupt change in the environment or a continuous, gradual change [33]. However, our approach is based on using two or more subsequent strikes, all of which are abrupt changes in the environment. Second, most existing models consider a single resistant variant (an exception is G. Martin et al. (2013) [31]), while the existing model for extinction therapy [16] works with a continuum of resistance effects. We choose to develop the simpler case in which we have discrete genetic variation. We, therefore, theoretically understand two-strike therapy as the prevention of evolutionary rescue.

### A.3 Derivation of extinction probabilities

Given the populations *S, R*_1_, *R*_2_ and *R*_1,2_ at time *τ*, we first compute the probability of no evolutionary rescue due to standing genetic variation. From evolutionary rescue theory (Appendix A.2), we know that the distribution of pre-existing rescue variants can be reasonably approximated by a Poisson distribution with a rate proportional to the number of rescue mutants at the beginning of the second strike (see Appendix A.4 for exceptions). This gives us a good estimate of the rescue probability due to *R*_2_ mutants. However, to estimate the rescue probability due to *R*_1,2_ mutants, we must consider the entire duration of the treatment because these cell types are resistant in both environments (*E*_1_ and *E*_2_). Therefore, in our model, the distribution of pre-existing *R*_2_ and *R*_1,2_ resistant cells is Poisson with a rate equal to 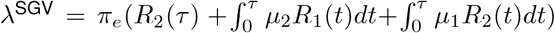, where *π*_e_ is the probability of establishment of a single resistant lineage and *µ*_1_ and *µ*_2_ are the rates of acquiring resistance to treatments 1 and 2. The probability of establishment depends on *b, d* and *c*(see Appendix A.5 for the derivation). Following the Poisson assumption, the probability that all pre-existing mutants go extinct in *E*_2_ is equal to 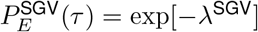.

To find the probability that no *de-novo* rescue mutants arise in *E*_2_, we assume that the generation of new mutants is a Poisson process, and the number of rescue mutants in *E*_2_ is Poisson distributed with a rate equal to 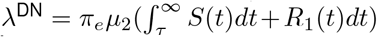. This rate is proportional to the rate of mutation from the sensitive cell fraction (*S* and *R*_1_) to the resistant cell fraction (*R*_2_ and *R*_1,2_) in *E*_2_.

The total extinction probability as a function of *τ* is given by the product of the probability of extinction of pre-existing and *de-novo* mutants,

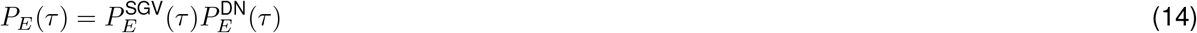

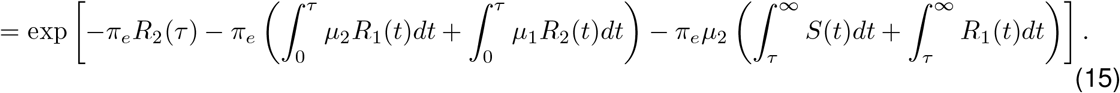

The first two terms inside the exponent account for the expected number of pre-existing *R*_2_ and *R*_1,2_ rescue mutants, and the last term accounts for the expected number of *de-novo* rescue mutants. To make the computation easier, we calculate the approximate values of the last two integral terms by integrating them until the population size is equal to one.

To compute the expression in (15) numerically, we evolve the deterministic logistic growth equations for dynamics in *E*_1_ (from *t* = 0 to *τ*, given in Figure 1) and in *E*_2_ (from *t* = *τ* till extinction, given below). In (16) and (17), we ignore the mutations to *R*_2_ and *R*_1,2_ and the changes in these resistant populations because we use only the deterministic decay of the sensitive population to calculate extinction probabilities.

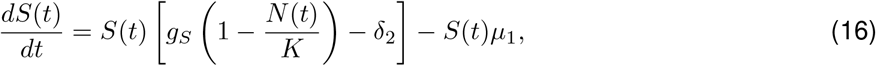

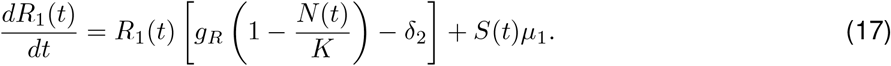

### A.4 Validity of the Poisson assumption

To compute extinction probabilities using our analytical model, we assume that the distribution of established rescue lineages (established lineages of *R*_1,2_ generated in either treatment phase and *R*_2_ in the second treatment) and *R*_2_ cells at the switching time is Poisson (rates given in Section A.3). This assumption is generally valid for the established rescue lineages, as they closely follow a pure Poisson point process, but it can break down for the pre-existing *R*_2_ cells in certain cases.

To investigate the distribution of *R*_2_ cells analytically, consider the stochastic master equation formulation of the Probability Mass Function (PMF) for the number of *R*_2_ cells 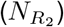 during treatment 1:

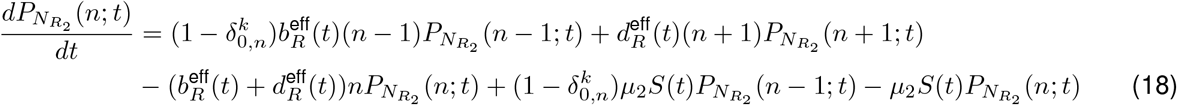

where *δ*_0,*n*_ denotes the Kronecker Delta function evaluated at (0, *n*) (*δ*_0,*n*_ = 1 if *n* = 0, and *δ*_0,*n*_ = 0 otherwise) and the *S* population is approximated as a deterministic function of time. This represents a birth-death-mutation process with time-inhomogenous event rates. Mutations from *R*_2_ to *R*_1,2_ are ignored. Following Getz [34], (18) can be converted to a single partial differential equation involving the probability generating function 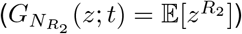:

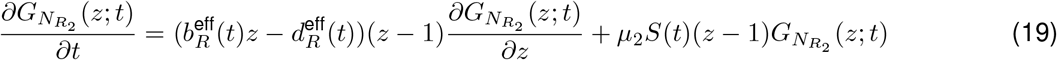

Equation (19) can be solved using the method of the characteristics [34]:

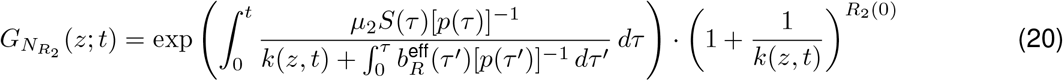

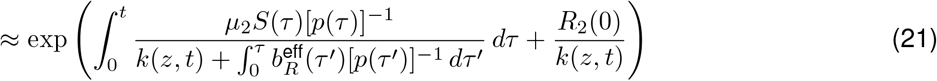

with 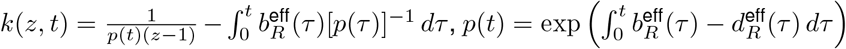 and with approximation (21) holding for *k*(*z, t*) large. There are two types of populations. Internal cells are *R*_2_ cells stemming from the initial *R*_2_(0) population and immigrant cells are *R*_2_ cells stemming from the mutant population. The random variables representing these populations are roughly independent at all times as the total *R*_2_ population contributes little to competition, so in (20) the first term represents the PGF of immigrant cells and the second term internal cells.

To obtain the probability of extinction of all *R*_2_ cells that arise before switching to the second treatment, let 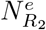 denote the number of *R*_2_ cells existing at the time of switching that eventually get established. Since 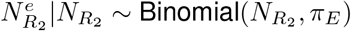:

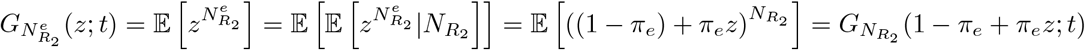

The probability of extinction from *R*_2_ cells arising before switching is just the mass of 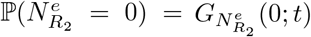. In the limit where the term 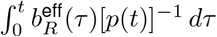 is negligible, which occurs when 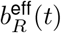 is made small at fixed 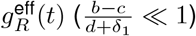 and using approximation (21), we get:

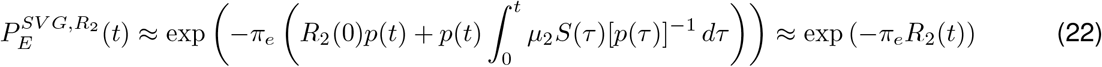

Expression (22) is what we use in (15). However when 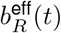 is made large at fixed 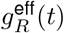 (or 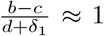), 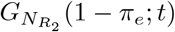 is strictly increased and can be made arbitrarily closer to 1^9^.

Consequently, the probability of extinction of the SGV *R*_2_ population calculated by our analytical model (Eq 2) is expected to be an *underestimate* of the probability of extinction of this population observed in our stochastic simulations especially when birth and death rates are comparable. For example, we observe this discrepancy in simulations with a zero cost of resistance and intermediate treatment dosages (second column of Figure 2 and Figure A.2). By SGV *R*_2_ population, we mean *R*_2_ cells descending from *R*_2_ cells initially present or from *R*_2_ cells arising from mutations of *S* cells during treatment 1.

### A.5 Derivation of the establishment probability

In this section, we provide the derivation of the establishment probability of a single lineage of resistant cells. This result is taken from Lambert, 2006 [35]. As described in Appendix A.2, the establishment probability is the probability with which resistant mutants establish themselves in the population and lead to evolutionary rescue. These mutants are then called rescue mutants.

We use the Continuous-time, real-valued Branching (CB) process to model the population dynamics of the resistant lineages. We chose this process because CB processes are the only diffusion processes that satisfy the additive property of the well-known BGW processes, which are commonly used to model stochastic population dynamics. Therefore, a CB process can be used to model the total population size summed over several lineages evolving independently. In our case, we use a CB process which is a continuous function of time, also called branching diffusion. As described in Lamperti, 1967 [36], branching diffusions are strong solutions of SDEs of the form:

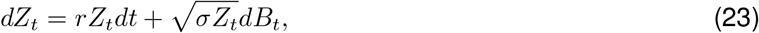

where *B* is the standard Brownian motion, *r* ∈ ℝ is the intrinsic growth rate, and *σ* ∈ ℝ^+^ is the reproduction variance, defined as [31]:

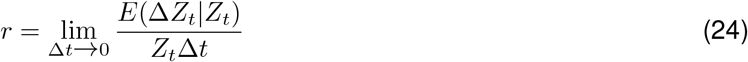

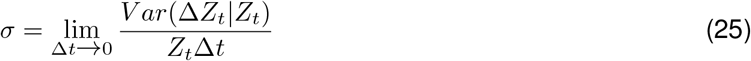

We use a general result from diffusion theory that the probability *u*(*z*_0_) of the diffusion hitting an absorbing barrier *z*_abs_, where *z*_0_ is the initial condition (*Z*_0_ = *z*_0_) is such that the function *u* solves the equation *Gu*(*z*) = 0 subject to appropriate boundary conditions. Here, *G* is the infinitesimal generator of the diffusion and characterizes the behaviour of the diffusion at small time intervals. For our one-dimensional diffusion (Equation 23), *G* is of the form,

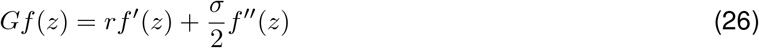

For the absorbing barrier *z*_abs_ = 0, we will obtain the extinction probability of a resistant population starting with *z*_0_ cells by solving *Gu*(*z*_0_) = 0, with boundary conditions *u*(*z*_abs_) = 1 and *u*(∞) = 0. The solution of this differential equation is,

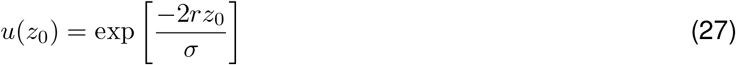

As explained in Martin et al., 2013 [31], the reproduction variance for a simple birth-death process with the birth rate *b* and death rate *d* can be approximated by *b* + *d*(turnover). Consequently, we obtain the following expression for the establishment probability, defined as one minus the extinction probability, for a resistant lineage starting from a single cell.

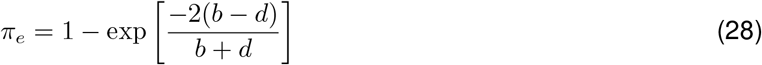

With a cost of resistance, we get,

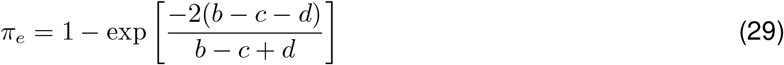

We use this result to compute the extinction probabilities with our deterministic analytical model in Section 2.1.

#### A.5.1 Alternate expression for the establishment probability

The Gillespie algorithm with birth rate *b* and death rate *d* is a numerical solution to the master stochastic equation:

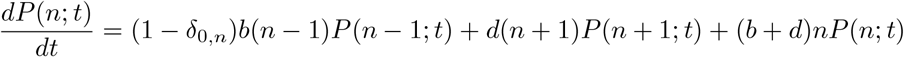

where *δ* is the Kronecker delta. Suppose we start with *X*_0_ = 1. This is a linear birth-death process, so the solution in terms of the probability-generating function *G* is [34]:

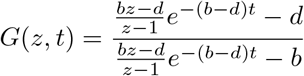

Note that *G*(0, *t*) = *P*(*X*_*t*_ = 0). To obtain the probability of extinction, the relevant event is ∃*t, X*_*t*_ = 0:

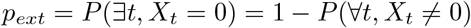

Now, we form a filtration 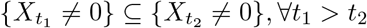, and thus:

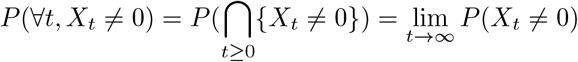

The limit 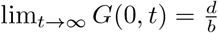, so 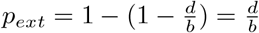. Therefore,

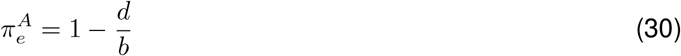

With a cost of resistance, *b* in (30) should be replaced with *b* − *c*. This expression is consistent with the method of simulation we use. The difference between the extinction probabilities obtained using (29) and (30) is negligible. To see this, firstly note that the first order Taylor approximations of *π*_*e*_(*b, d*) and 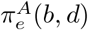 centered about (*q*_0_, *q*_0_) are 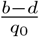 and so behave quite similarly for small *g* = *b*−*d*. Their values are made most different as (*b, d*) → (∞, *d*) for any fixed *d* and in this limit become *π*_*e*_ → 1 − *e*^−2^ and 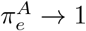. These values differ by ≈ 0.13, which is quite large, but the more relevant difference is between expressions of the form 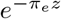 and 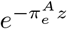. As (*b, d*) → (∞, *d*), the maximum difference is around 0.05 and is obtained at *z* ≈ 1.07. Finally, any qualitative comparisons between probabilities of extinction remain the same independent of choice of *π*_*e*_ or 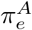.

### A.6 Stochastic Simulation Model

The total population at time *t* is *N*(*t*) = *S*(*t*) + *R*_1_(*t*) + *R*_2_(*t*) + *R*_1,2_(*t*), as defined in Section 2.1. The code developed for the implementation of this model is designed to be highly versatile and easy to modify. It can be extended to analyse more complex systems with more treatments and corresponding cell types.

#### A.6.1 Gillespie-like Implementation

The stochastic simulation algorithm (SSA), commonly referred to as the Gillespie algorithm [37], is a Monte Carlo method initially devised in 1977 for modelling the temporal evolution of chemical reactions in a well-mixed system. We implement a version of the Gillespie algorithm to simulate the population dynamics of our system in the context of two-strike therapy. The idea is that given a set of rates corresponding to birth, death and mutation events, we can track the size of each subpopulation. These rates are specified for all cell types and can change with time or in response to the environment. The basic steps of our algorithm are as follows:

1. Initialize the system by specifying an initial population for all cell types (*S*(0), *R*_1_(0), *R*_2_(0), *R*_1,2_(0)) and setting the time to zero. Define all possible demographic events and their corresponding rates.
2. Calculate the rate of any one event occurring, which is the sum of all individual rates (denoted by *ω*_*i*_(*t*) for each event *i*). Then, obtain the time interval after which the next event will take place. To do this, generate a sample from an exponential random variable with the rate parameter equal to the total rate Σ_*i*_ *ω*_*i*_(*t*). Alternatively, one can generate a random number *z*_1_ from the uniform distribution between 0 and 1 and use the following formula to determine the time interval for the next event: *t*_int_ = − ln(*z*_1_)*/* Σ_*i*_ *ω*_*i*_(*t*)
3. Calculate event probabilities for the next event by dividing the individual event rates by the total rate: *p*_*i*_(*t*) = *ω*_*i*_(*t*)*/* Σ _*i*_ *ω*_*i*_(*t*). Use these probabilities to select the next event by generating a random number *z*_2_ between 0 and 1. The chosen event *k* would be the largest *j* such that 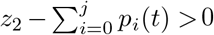.
4. Implement the chosen event by updating the population of the corresponding cell types. Increment time *t* = *t* + *t*_int_. For example, if the chosen event is the birth of an *R*_2_ cell, then *R*_2_(*t* + *t*_int_) = *R*_2_(*t*) + 1.
5. Repeat steps 2-4 till a stopping condition is reached.

**Table A.1:** Stopping conditions and corresponding outcomes of a simulation. In the second condition, *T* is the maximum simulation time defined at the beginning. If we see a significant number of outcomes with persistence (more than 10%), it means that *T* is not large enough and the simulation is run again with a higher *T* value. In the third condition, the threshold 0.99*K* is arbitrary. The outcome remains the same as long as the threshold is close to *K*. We take the minimum of two quantities for cases where *K* is much larger than the initial population size, and a threshold of *N*_max_ is enough to declare the outcome. In all cases, *K* ≥ *N*(0), and *N*_max_ > *N*(0), so persistence in the last condition is not observed. However, the condition is mentioned for the sake of completion.

| Stopping condition | Outcome condition | Outcome |
| --- | --- | --- |
| $N(t) = 0$ | $N(t) = 0$ | Extinction |
| $t \geq T$ | $N(t) \geq N(0)$<br>$N(t) < N(0)$ | Progression<br>Persistence |
| $N(t) \geq \min(0.99K, N_{\max})$ | $N(t) \geq N(0)$<br>$N(t) < N(0)$ | Progression<br>Persistence |

Note that the effects of carrying capacity and treatment are included while specifying birth and death rates in Section A.6.2. Simulations are stopped under one of three conditions: if the population goes extinct, if it exceeds the maximum simulation time, or if it exceeds some maximum population size. Similarly, the outcome of one run of a simulation can be one of three possibilities: extinction (*N*(*t*) = 0), progression (*N*(*t*) ≥ *N*(0)) or persistence (0 *< N*(*t*) *< N*(0)). Note that there is not a one-to-one correspondence between stopping conditions and simulation outcome (see Table A.1)

#### A.6.2 Determining rates of demographic event

Following our variant of the Gillespie algorithm, one must define all possible demographic events at the beginning of the simulation and define rates corresponding to those events at each time step. Note that an individual event includes the type of event and the type of cell. For example, a mutation event *S* → *R*_1_ is one individual event and has a rate specified for it. Similarly, the birth of an *S* cell is a different event than the birth of an *R*_1_ cell. All simulation parameters used to compute the individual event rates are listed in Table 1.

We derive the birth and death rates for all cell types from the deterministic Logistic model for population growth.

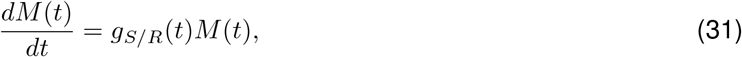

where *g*_*M*_ (*t*) is the growth rate of subpopulation *M* ∈ {*S, R*_1_, *R*_2_, *R*_1,2_}. The growth rates of sensitive and resistant subpopulations differ by the cost of resistance,

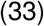

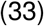

where the total treatment-induced death rate in environment *i* is *δ*_*i*_, and the presence of this term depends on the sensitivity of different cell types in both environments. For example, the growth rate of *R*_1_ cells in *E*_1_ will not include the treatment-induced death term, but in *E*_2_ will have a *δ*_2_ term.

We can write the intrinsic growth rates as the difference between intrinsic birth and death rates like so:

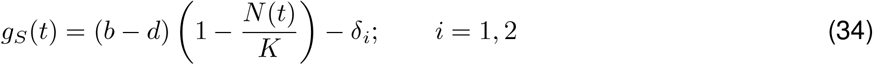

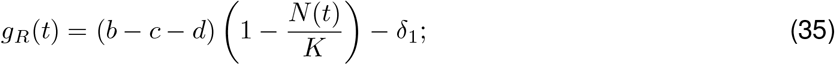

where *b, d* and *c* are the intrinsic birth rate, death rate and the cost of resistance. Now, we separate the positive and negative parts of the subpopulation growth rates to find the effective birth and death rates to use for our simulations.

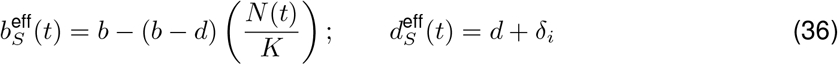

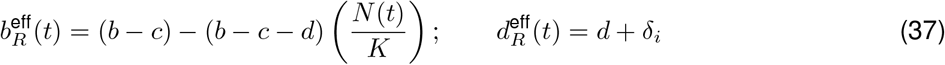

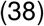

We use Equations 36 and 37 in our simulations. Note that the separation of positive and negative terms in the last step is not unique. The only constraint is that the separation follows the condition required for the implementation of carrying capacity, i.e. when the total population is equal to *K*, we must have 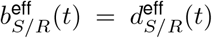 (without treatment). We chose a simple way to implement this by taking a density-dependent birth rate and a density-independent death rate. Additionally, it is convenient and intuitive to group the death-rate terms together, like we do in the 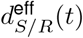.

For mutation events, we consider the same mutation rate 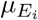 for all cell types, which may change with the environment. Once the source population for an event is chosen (say, *S* cells), the target cell is chosen according to the rates of acquiring each type of mutation. These mutation probabilities from one cell type to the other can be modified.

#### A.6.3 Switching to the second treatment

For this study, we consider two environments *E*_1_ and *E*_2_, each corresponding to the two strikes or treatments. Now we ask the question: how does the environment change with time? In other words, what is the condition under which we switch treatments and apply the second strike? Two-strike therapy aims to minimize the probability of evolutionary rescue, and the expected number of rescue mutants depends on the population size and time at which we switch between treatments. The population size at the time of switching is *N*(*τ*).

It is necessary to understand the behaviour of this variable *N*(*τ*) in order to evaluate the efficacy and limitations of two-strike therapy. As reasoned by Gatenby et al. (2020) [16]and supported by our results, the hypothesis is that the optimal *N*(*τ*) would be close to *N*_min_. To test this, we run a set of simulations with the same parameter values but with different random seeds. For a single random seed, we run multiple simulations (using the algorithm in Appendix A.6.1), each with a different *N*(*τ*). To produce Figures 2(A) and A.2, we run simulations with 100 random seeds for a single set of parameters and then calculate extinction probabilities for various values of *N*(*τ*). The algorithm is as follows:

1. Select a set of parameter values. These parameters are kept constant throughout.
2. Set random seed for this set of simulations, thus eliminating differences due to stochasticity.
3. Run the first simulation without any second strike. Equivalently, set *N*(*τ*) equal to zero for this run. From the results of this run, save the values of *N*_min_ and its corresponding time point *t*(*N*_min_).
4. Run the remaining set of simulations with increasing values of *N*(*τ*). Record the outcomes of all the runs (according to Table A.1. See Figure 2(B) for an illustration.
5. Repeat steps 2-4 for the desired number of random seeds. Each set of simulations with a different random seed is independent and will have different values of *N*_min_. However, the values of *N*(*τ*) will remain the same for each independent realisation in order to calculate extinction probabilities.
6. Calculate extinction probabilities using outcomes for each random seed corresponding to the values of *N*(*τ*).

### A.7 Choosing the default parameter values

The default parameter set used for the stochastic simulations and numerical solutions are listed in Table 1. The carrying capacity is set equal to the total initial population size for consistency with previous studies on extinction therapy [16]. Moreover, we show that changing the carrying capacity has negligible effects on our results (Section 3.8). The default cost of resistance is set to a high level (half of the intrinsic birth rate), but the case with zero cost of resistance is also studied in detail to elucidate the effect of this parameter (Section 3.3). The size of the initial resistant populations is also set to 10^−4^ times the initial sensitive population, which is higher than what we obtain using the growth model described in Appendix A.1. However, in our results, we consider the case with *R*_2_= 0. Moreover, our default parameter value shows that two-strike therapy can work with a conservative estimate of the initial resistant population.

### A.8 Metrics to compare treatment outcomes using output from the analytical model

After devising a way to capture the drop in extinction probabilities as *N*(*τ*) increases, we wanted to compare these curves for different sets of parameters to determine the optimal conditions for two-strike therapy. After observing the (nearly) monotonously decreasing nature of the *N*_*q*_ vs *q* plot, we came up with three metrics to achieve this goal. The first metric is the normalised area under the *N*_*q*_ vs *q* curve. A larger area under the curve (AUC) can mean one of two things – first, that the drop in extinction probability occurs at a larger *N*(*τ*) or second, that the drop occurs gradually over a larger range of *N*(*τ*) values. The first scenario is favourable for ET, and the second one can be beneficial in some cases. If the drop is close to the initial population, then it is easier to implement ET, and if the drop is gradual, then there are chances to obtain high extinction probabilities at high *N*(*τ*) values. Clearly, this metric is not very accurate, but it is a reasonable choice considering the general trend of *N*_*q*_ vs *q* curves showing a sudden drop in extinction probabilities. Therefore, the set of parameters for which the area under the *N*_*q*_ curve is higher is expected to result in a better outcome in terms of ease of implementation, higher extinction probabilities, or both.

The second metric we propose allows us to find the population size at which the drop in extinction probability occurs. We define the critical second strike threshold as *N*_*c*_ =mean(*N*_0.1_, *N*_0.9_). The higher the *N*_*c*_ for a system, the easier it will be to implement two-strike therapy. This metric has the advantage of approximating the time of the drop, but it is accurate only if the drop is from a value as high as 0.9 to a value as low as 0.1. Other values for the definition of *N*_*c*_ may be considered, but it is hard to find appropriate values that work for a wide range of parameter sets. The third metric is similar to the second one and uses the *N*_0.5_*/N*(0) as a measure of the population size at which this drop occurs.

We compare the three metrics in Figure A.1 and see that they produce similar results. Furthermore, these results are consistent with the trends observed when comparing regions of high-*P*_*E*_ (Figure 4(B)).

### A.9 Supplementary Tables and Figures

**Figure A.1:**
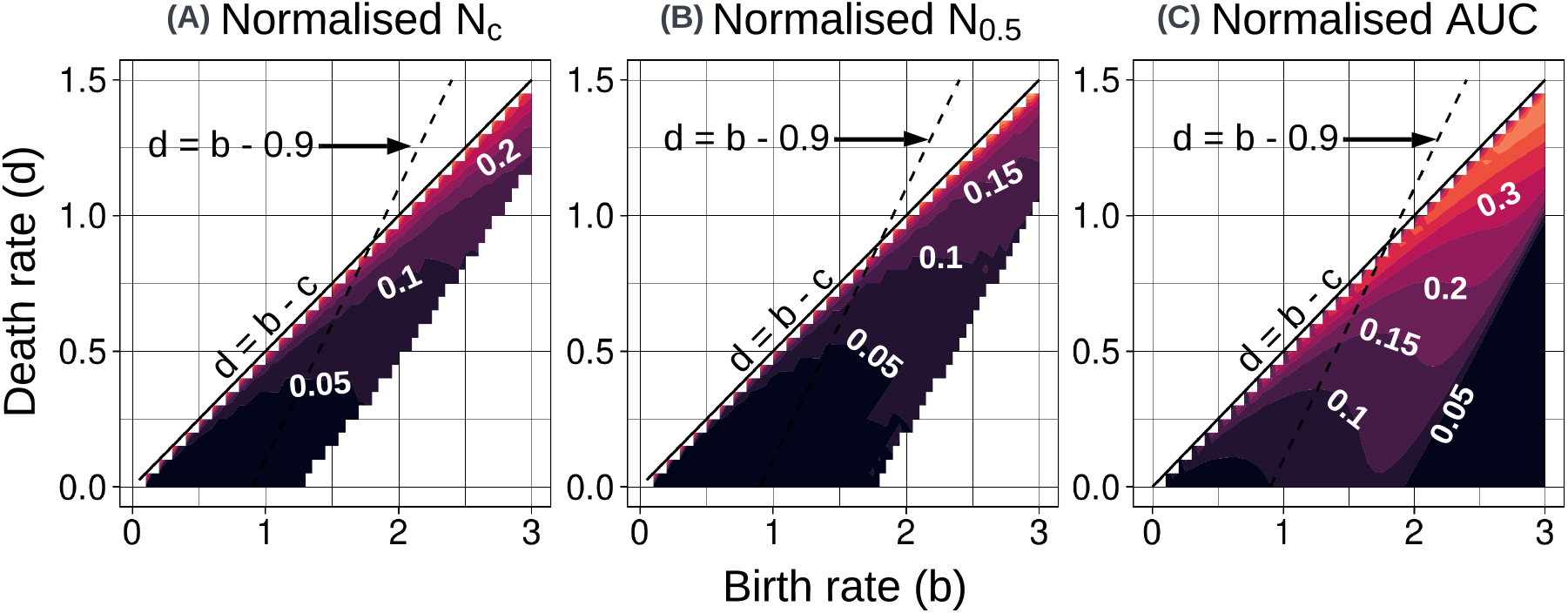
Comparing treatment outcomes for different parameter values in the *b* −*d* space using three different metrics. For each metric, values of birth and death rates are chosen such that the resistant cells have a positive growth rate. The dashed line indicates the set of birth and death rates which result in a constant intrinsic growth rate (equal to the default value of *g*_*S*_ = 0.9).**(A)** The metric is defined as the normalised mean of *N*_0.1_ and *N*_0.9_. The bottom right region gives invalid values using this metric because *N*_0.9_ is not defined for those parameters. This is because the optimal extinction probabilities in that region are lower than 0.9. **(B)** The metric is defined as the normalised *N*_0.5_. Similar to the metric used in plot A, the parameters in the bottom right region give optimal extinction probabilities less than 0.5. **(C)** The normalised area under the curve of the *N*_*q*_ vs *q* plot for parameters in the *b* − *d* space. This metric is favoured because it gives smooth curves on the plot, which also does not contain an invalid region.

**Figure A.2:**
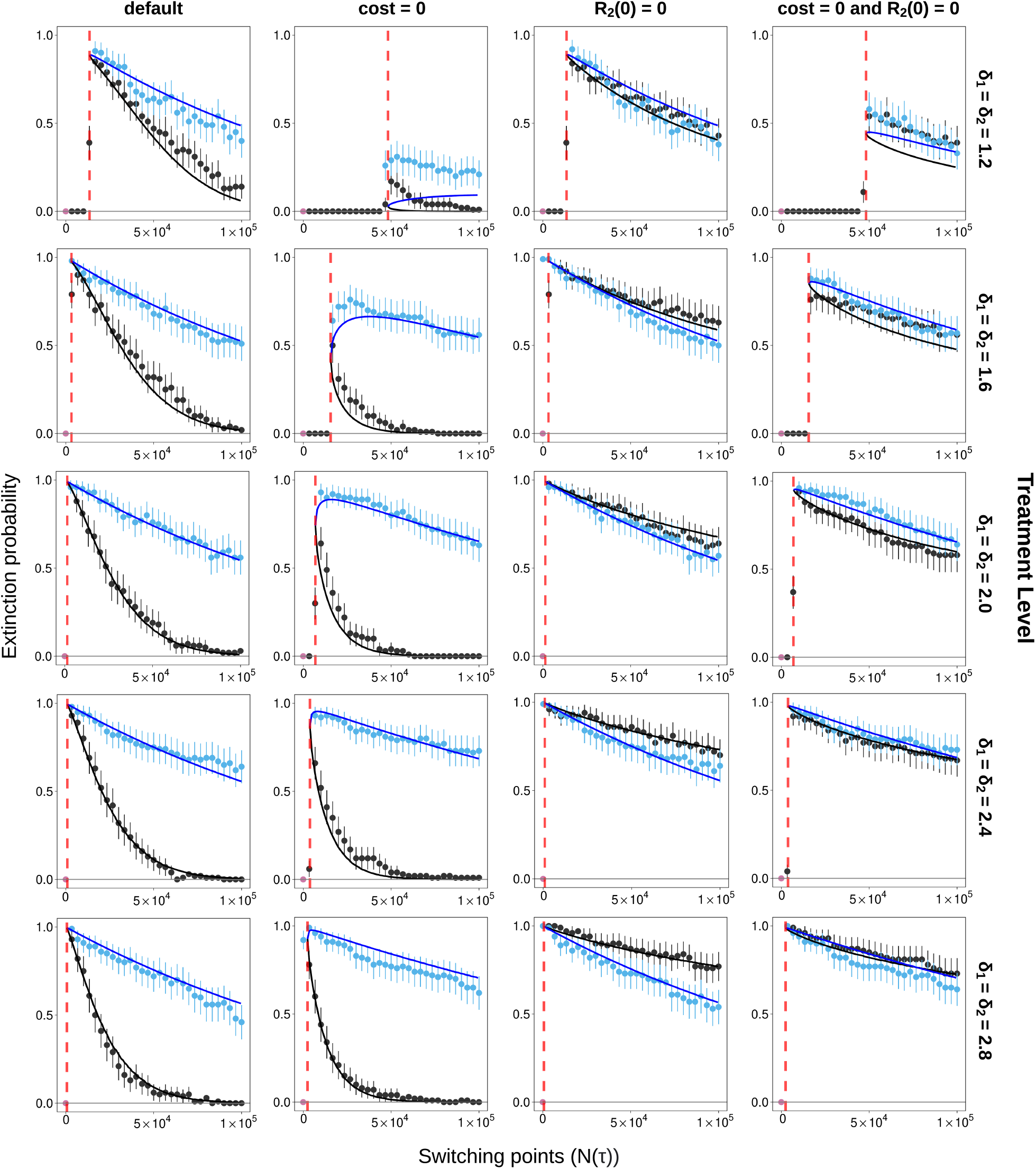
Simulation results for points of switching before and after *N*_min_. For the same random seed, extinction probabilities for different switching points (see Appendix A.6) are plotted. The black points indicate extinction probabilities when the points of switching are before the *N*_min_, and blue points indicate *N*(*τ*)’s implemented after *N*_min_. Pink dots show the extinction probability in the absence of a second treatment. See Figure 2(B) for the legend. All parameters except the cost of resistance, treatment level and initial *R*_2_ population are set to their default values. Error bars show 95% binomial proportion confidence intervals. All extinction probabilities are obtained by considering the outcomes of 100 runs of the simulation with the different random seeds.

**Figure A.3:**
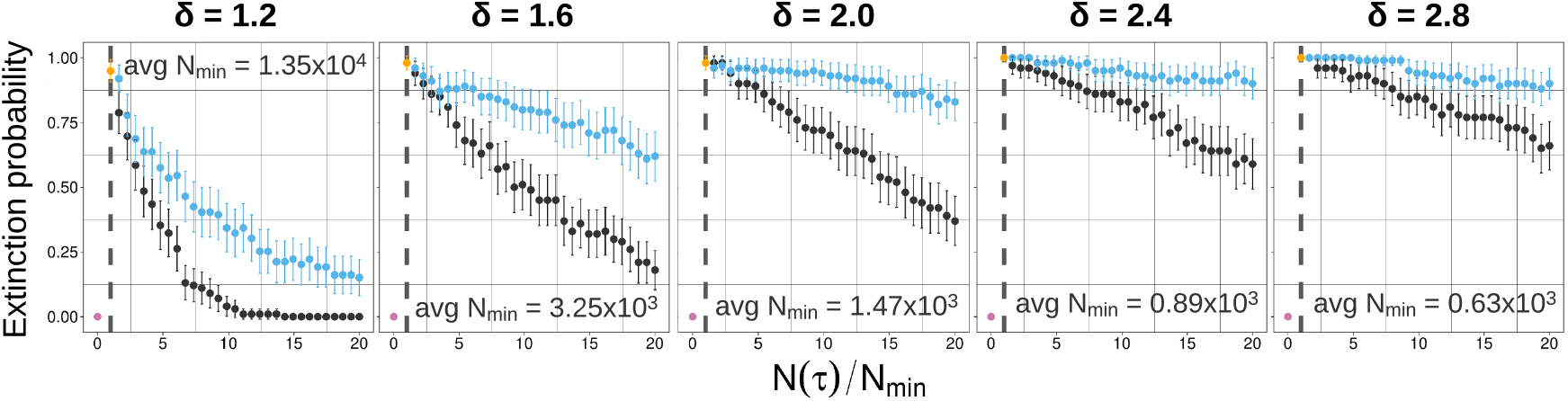
Extinction probabilities for several switching points relative to the *N*_min_ (different for each of 100 independent runs). Black(blue) dots show simulation results for before(after)-nadir switching points.

**Figure A.4:**
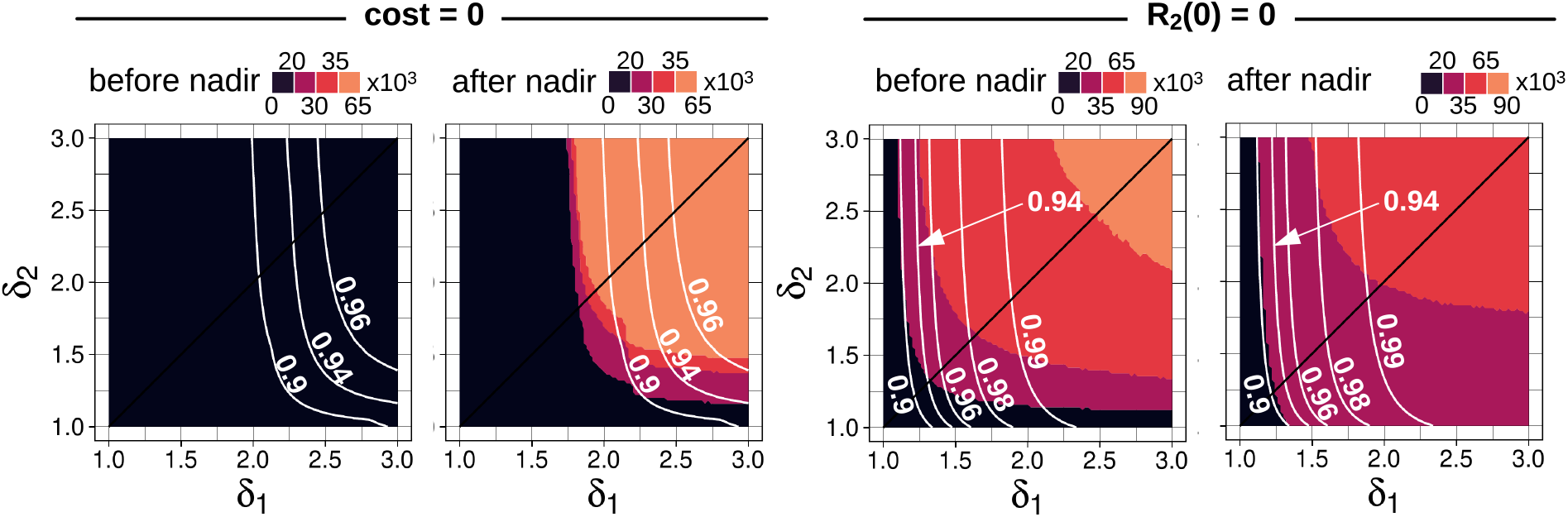
Heatmaps (obtained from the analytical model) showing the range of *N*(*τ*) values that give a high extinction probability (≥ 0.8) for different combinations of treatment levels *δ*_1_ and *δ*_2_. The case with cost= 0 is shown on the left, and the case with *R*_2_(0) = 0 is on the right. For each case, both before-nadir and after-nadir switching points are considered. White lines indicate optimal extinction probability contours. In the leftmost panel, no high-*P*_*E*_ regions exist.

**Figure A.5:**
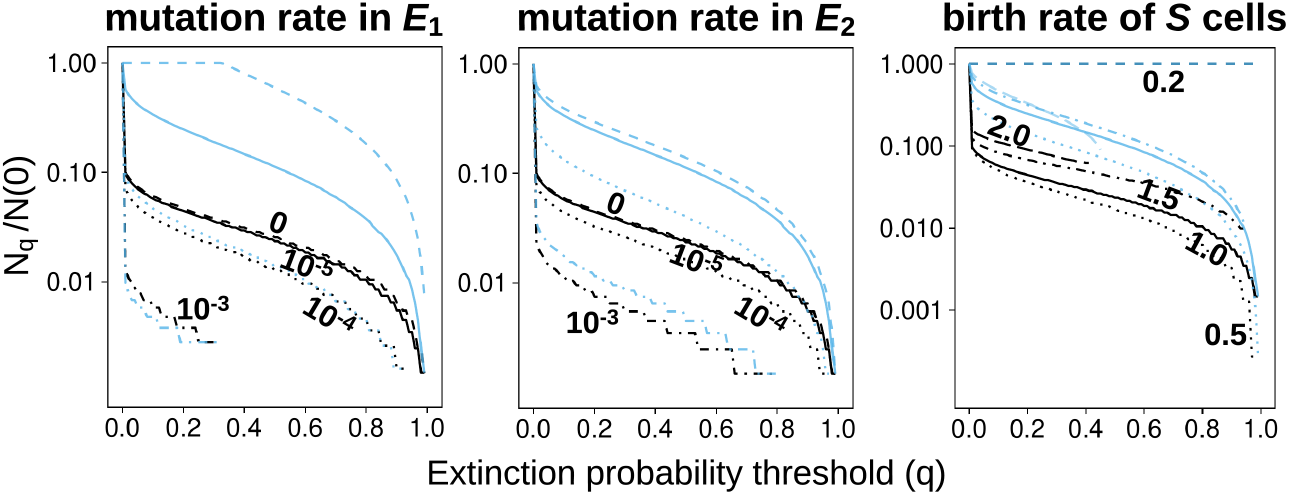
Normalised *N*_*q*_ vs *q* plot for different values of total mutation rates in *E*_1_ and *E*_2_, and the intrinsic birth rate. The value of 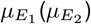 is kept constant at 10^−5^ when 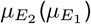 is varied. Changing the total mutation rate in both environments individually has the same effect (qualitatively). A higher mutation rate results in lower extinction probabilities. This figure is obtained using the analytical model only.

**Figure A.6:**
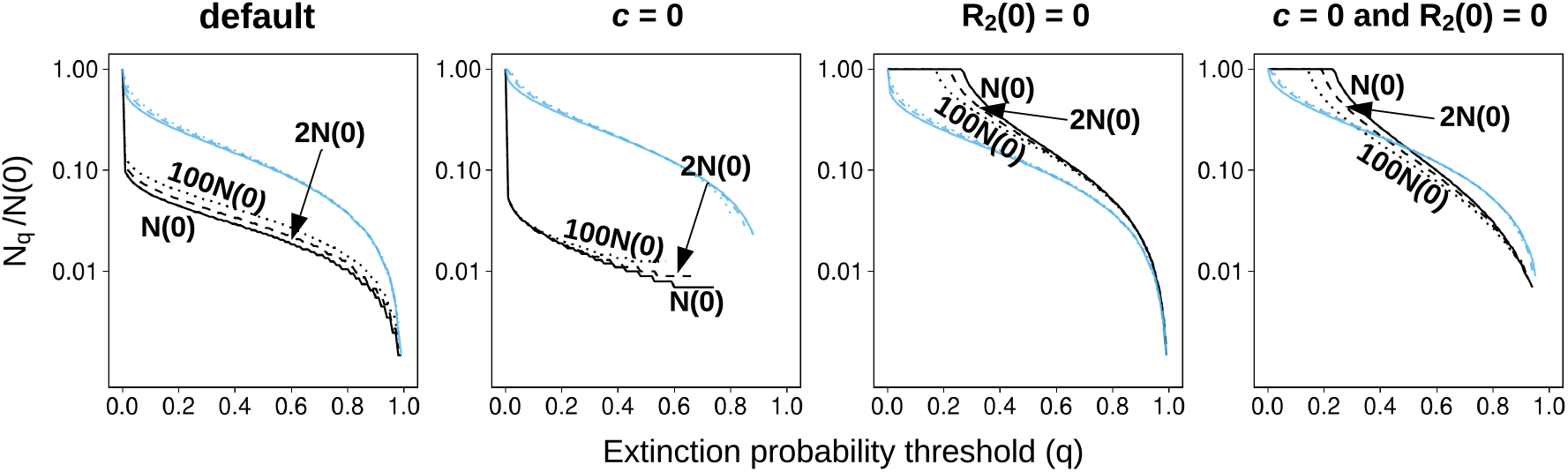
Normalised *N*_*q*_ vs *q* plot, obtained using the analytical mode, for different values of the carrying capacity under four conditions – default, no cost of resistance, no initial *R*_2_ population, and the case with *c* = 0 and *R*_2_(0) = 0. The last case reveals the effect of changing the carrying capacity in isolation. Black(blue) lines indicate before(after)-nadir switching points. The line type (solid, mixed, dotted) is the same for both before and after nadir curves.

**Figure A.7:**
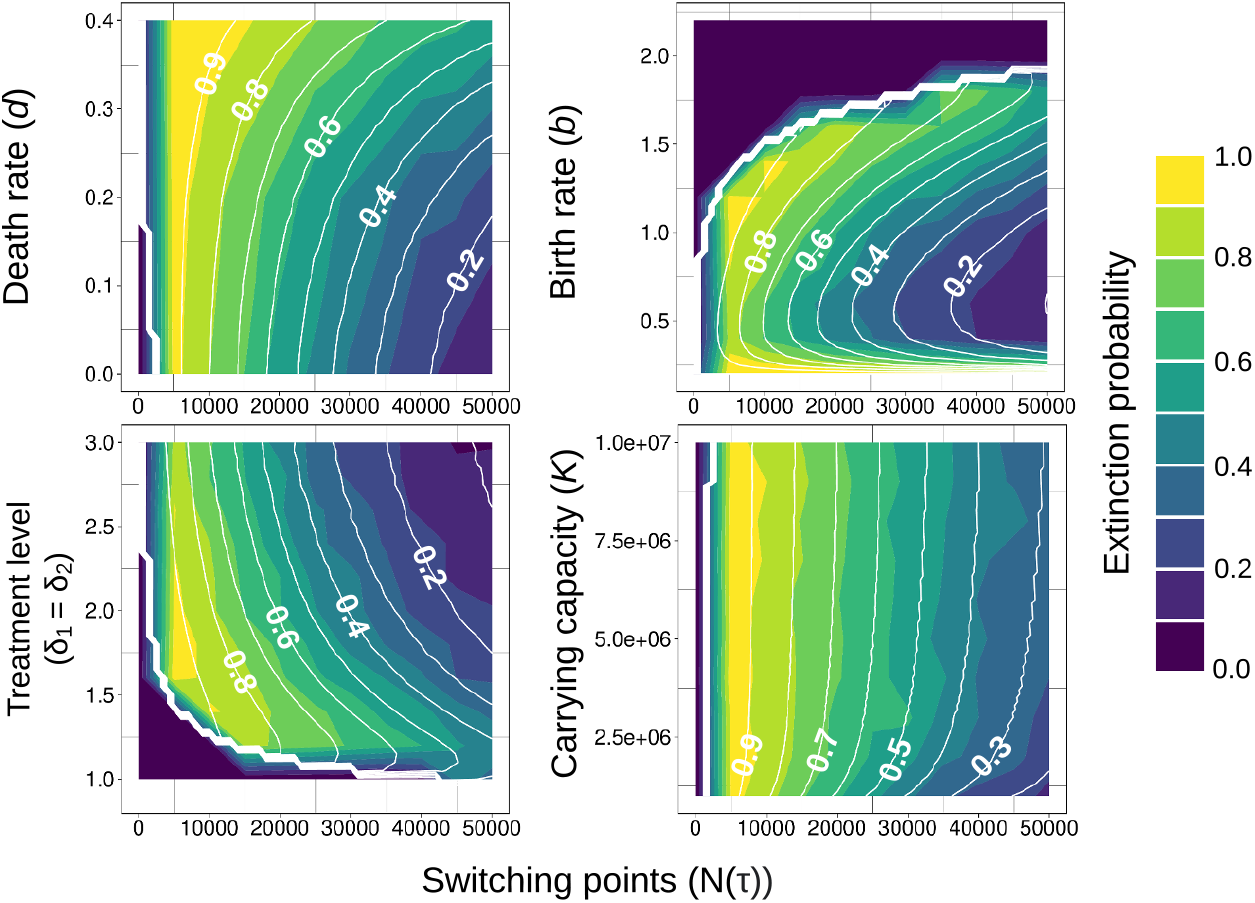
Extinction probability heatmaps for four system parameters. All other parameters are from the default parameter set (Table 1). In all the heatmaps, the solid white contours (with labels) show analytical results. Stochastic simulation results are denoted by the colour scale. Extinction probabilities from the stochastic model are obtained by using the outcomes of 500 simulations with the same parameter values and initial conditions. In the bottom-left panel, treatment levels for both environments are altered together (*δ*_1_=*δ*_2_). The default treatment level is *δ*= 2.0 per unit of time. We do not consider treatment levels below 0.9 because that is the intrinsic growth rate of *S* cells, due to which *δ<* 0.9 will only give positive growth rates for all cells in the population. The dark regions in the plots (e.g. top-left region in the top-right panel) have low extinction probabilities because *N*(*τ*)*<N*_min_ at those points.

**Figure A.8:**
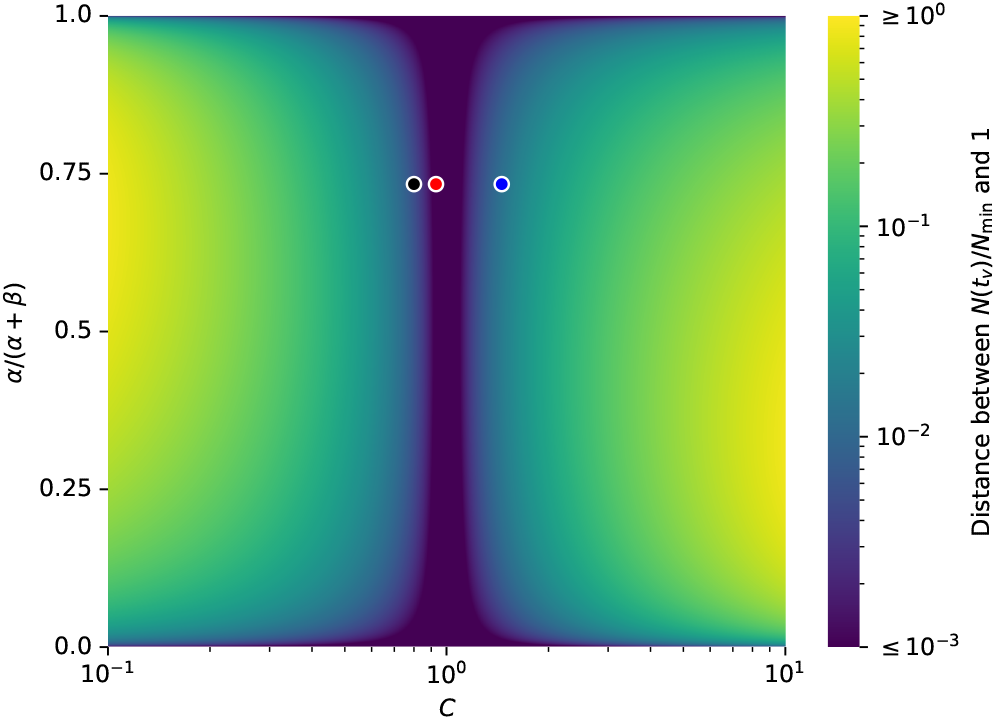
Measuring how close 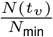 (calculated using Lemma A.1) is to 1, where *t*_*v*_ can stand for the time of minimal standing genetic variants (SGV), time of minimal de-novo variants (DN), or time of optimal probability of extinction (opt). *α* and *β* were defined in section A.1.2 to be *δ*_1_ − *γ*_*S*_ and *γ*_1_, respectively. *C* is an axis depending on variables *δ*_2_, *γ*_1_, *c*_*i*_, *π*_*i*_ and its functional form is set to 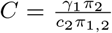 for *v* = SGV, 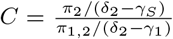 or *v* = DN, and 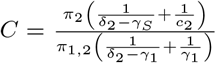 for *v* = opt (see sections A.1.2.3, A.1.2.2, and A.1.2.4 respectively). The black, blue, and red dots indicate the default parameter choices for *v* = SGV, DN, opt, respectively.

## Footnotes

1 If *γ*_2_ = *γ*_*S*_, that is, *c*_2_ = 0, then the term 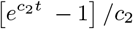 in the expression for *R*_2_(*t*) should be replaced by *t*, or equivalently by 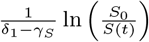.

2 The quantity 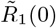 represents the initial *R*_1_ population that would generate the same long-run *R*_1_ population in the absence of mutations from *S* cells to *R*_1_. It is positive. The quantity 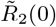 represents the additional initial *R*_2_ population compared to a reference case detailed below. It may be positive or negative.

3 In the absence of a cost of resistance: *c*_*i*_ = 0, the previous formula becomes 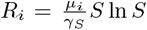, and with our other default parameter values, we obtain *R*_*i*_(0) ≃ 38, still lower than 100.

4 With our default parameter values, 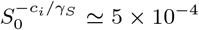, and the approximation in (11) is excellent. It is less good for small costs of resistance, e.g., if *S*_0_ = 10^6^ and *c*_*i*_*/γ*_*S*_ = 1*/*6, then 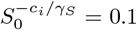 and we make a 10% error.

5 For instance, slightly lowering the resistance costs *c*_1_ or *c*_2_ would increase the ratio *α*_*S*_*/α*_1_, and for *c*_1_ = *c*_2_ = 0.47, we already obtain that *t*_*opt*_ *> t*_*min*_, i.e. we should now switch after rather than before the nadir.

6 In the reference case discussed above, it may be shown that for any Δ *>* 0, *RM*(*t*_*opt*_ + Δ) *≤ RM*(*t*_*opt*_ − Δ).

7 The situation would be more complex in a model with competition, as reducing the sensitive population quickly leads to the competitive release of the resistant populations. Thus, a larger value of *δ*_1_ would also have negative sides, and the net effect should be estimated more carefully.

8 This effect would disappear if the number of mutations from *R*_1_ to *R*_1,2_ was assumed proportional not only to *R*_1_ but also to its net growth-rate. The effect would be partially restored if the number of mutations was assumed proportional to the net birth rate.

9 Intuitively, as the intrinsic birth rate is made more comparable to the intrinsic death rate at some growth rate, the *R*_2_ population follows a more heavy-tailed distribution than what a corresponding Poisson approximation would suggest. This leaves more mass at the 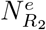 = 0 absorbing barrier, and hence a higher probability of extinction

## References

[1] Yoh Iwasa, Martin A Nowak, and Franziska Michor. “Evolution of Resistance During Clonal Expansion”. In: Genetics 172.4 (Apr. 2006), pp. 2557–2566. ISSN: 1943-2631.

[2] Mariyah Pressley et al. “Evolutionary Dynamics of Treatment-Induced Resistance in Cancer Informs Understanding of Rapid Evolution in Natural Systems”. In: Frontiers in Ecology and Evolution 9 (2021). ISSN: 2296-701X. URL: https://www.frontiersin.org/articles/10.3389/fevo.2021.681121 (visited on 03/30/2023).

[3] Mel Greaves and Carlo C. Maley. “Clonal evolution in cancer”. en. In: Nature 481.7381 (Jan. 2012). Number: 7381 Publisher: Nature Publishing Group, pp. 306–313. ISSN: 1476-4687. DOI: 10.1038/nature10762. URL: https://www.nature.com/articles/nature10762 (visited on 09/01/2022).

[4] Kirill S. Korolev, Joao B. Xavier, and Jeff Gore. “Turning ecology and evolution against cancer”. en. In: Nature Reviews Cancer 14.5 (May 2014), pp. 371–380. ISSN: 1474-1768.

[5] Pedro M. Enriquez-Navas, Jonathan W. Wojtkowiak, and Robert A. Gatenby. “Application of Evolutionary Principles to Cancer Therapy”. In: Cancer Research 75.22 (Nov. 2015), pp. 4675–4680. ISSN: 0008-5472. DOI: 10.1158/0008-5472.CAN-15-1337. eprint: https://aacrjournals.org/cancerres/article-pdf/75/22/4675/2937393/4675.pdf. URL: https://doi.org/10.1158/0008-5472.CAN-15-1337.

[6] C. Athena Aktipis et al. “Overlooking Evolution: A Systematic Analysis of Cancer Relapse and Therapeutic Resistance Research”. en. In: PLOS ONE 6.11 (Nov. 2011), e26100. ISSN: 1932-6203.

[7] Jeffrey West et al. “A survey of open questions in adaptive therapy: Bridging mathematics and clinical translation”. In: eLife 12 (Mar. 2023). Ed. by Richard M White. Publisher: eLife Sciences Publications, Ltd, e84263. ISSN: 2050-084X. DOI: 10.7554/eLife.84263. URL: https://doi.org/10.7554/eLife.84263 (visited on 03/30/2023).

[8] Helen C. Monro and Eamonn A. Gaffney. “Modelling chemotherapy resistance in palliation and failed cure”. In: Journal of Theoretical Biology 257.2 (2009), pp. 292–302. ISSN: 0022-5193. DOI: 10.1016/j.jtbi.2008.12.006. URL: https://www.sciencedirect.com/science/article/pii/S0022519308006334.

[9] Robert A. Gatenby et al. “Adaptive therapy”. In: Cancer Research 69.11 (June 2009), pp. 4894–4903. ISSN: 00085472. DOI: 10.1158/0008-5472.CAN-08-3658.

[10] Helen K. Alexander et al. “Evolutionary rescue: linking theory for conservation and medicine”. en. In: Evolutionary Applications 7.10 (2014), pp. 1161–1179. ISSN: 1752-4571.

[11] Robert A. Gatenby, Jingsong Zhang, and Joel S. Brown. “First strike-second strike strategies in metastatic cancer: Lessons from the evolutionary dynamics of extinction”. In: Cancer Research 79.13 (2019), pp. 3174–3177. ISSN: 15387445.

[12] Brian Dennis et al. “Allee effects and resilience in stochastic populations”. en. In: Theoretical Ecology 9.3 (Sept. 2016), pp. 323–335. ISSN: 1874-1746.

[13] Robert A. Gatenby and Joel S. Brown. “Integrating evolutionary dynamics into cancer therapy”. en. In: Nature Reviews Clinical Oncology 17.11 (Nov. 2020). Number: 11 Publisher: Nature Publishing Group, pp. 675–686. ISSN: 1759-4782. DOI: 10.1038/s41571-020-0411-1. URL: https://www.nature.com/articles/s41571-020-0411-1 (visited on 03/30/2023).

[14] Christophe Meille et al. “Revisiting Dosing Regimen Using Pharmacokinetic/Pharmacodynamic Mathematical Modeling: Densification and Intensification of Combination Cancer Therapy”. en. In: Clinical Pharmacokinetics 55.8 (Aug. 2016), pp. 1015–1025. ISSN: 0312-5963, 1179-1926.

[15] Shaon Chakrabarti and Franziska Michor. “Pharmacokinetics and Drug Interactions Determine Optimum Combination Strategies in Computational Models of Cancer Evolution”. en. In: Cancer Research 77.14 (July 2017), pp. 3908–3921. ISSN: 0008-5472, 1538-7445.

[16] Robert A. Gatenby et al. “Eradicating metastatic cancer and the eco-evolutionary dynamics of Anthropocene extinctions”. In: Cancer Research 80.3 (Feb. 2020), pp. 613–623. ISSN: 15387445.

[17] NCT04388839: Evolutionary Inspired Therapy for Newly Diagnosed, Metastatic, Fusion Positive Rhabdomyosarcoma. May 2020. URL: https://clinicaltrials.gov/study/NCT04388839 (visited on 09/27/2020).

[18] NCT05189457: A Phase IIA Study of Sequential (“First Strike, Second Strike”) Therapies, Modeled on Evolutionary Dynamics of Anthropocene Extinctions, for High Risk Metastatic Castration Sensitive Prostate Cancer. Dec. 2021. URL: https://clinicaltrials.gov/study/NCT05189457 (visited on 01/25/2022).

[19] NCT06409390: A Pilot Study of Sequential (“First Strike, Second Strike”) Therapies, Modeled on Evolutionary Dynamics of Anthropocene Extinctions, for Hormone Positive Metastatic Breast Cancer. May 2024. URL: https://clinicaltrials.gov/study/NCT06409390 (visited on 04/26/2024).

[20] Yannick Viossat and Robert Noble. “A theoretical analysis of tumour containment”. In: Nature Ecology and Evolution 5.6 (June 2021), pp. 826–835. ISSN: 2397334X.

[21] Maximilian Strobl et al. Turnover modulates the need for a cost of resistance in adaptive therapy. en. Pages: 2020.01.22.914366 Section: New Results. Mar. 2020. DOI: 10.1101/2020.01.22.914366. URL: https://www.biorxiv.org/content/10.1101/2020.01.22.914366v2 (visited on 03/30/2023).

[22] Damon R. Reed et al. “An evolutionary framework for treating pediatric sarcomas”. In: Cancer 126.11 (2020). eprint: https://onlinelibrary.wiley.com/doi/pdf/10.1002/cncr.32777, pp. 2577–2587. ISSN: 1097-0142. DOI: 10.1002/cncr.32777. URL: https://onlinelibrary.wiley.com/doi/abs/10.1002/cncr.32777 (visited on 03/31/2023).

[23] Seth I. Felder, Jason B. Fleming, and Robert A. Gatenby. “Treatment-induced evolutionary dynamics in nonmetastatic locally advanced rectal adenocarcinoma”. en. In: Advances in Cancer Research. Vol. 151. Elsevier, 2021, pp. 39–67. ISBN: 978-0-12-824078-6. DOI: 10.1016/bs.acr.2021.02.003. URL: https://linkinghub.elsevier.com/retrieve/pii/S0065230X21000191 (visited on 01/10/2023).

[24] J H Goldie, A J Coldman, and G A Gudauskas. “Rationale for the Use of Alternating Non-Cross - Resistant Chemotherapy”. en. In: Cancer Treatment Reports 66 (1982).

[25] A.J. Coldman and J.H. Goldie. “A model for the resistance of tumor cells to cancer chemotherapeutic agents”. en. In: Mathematical Biosciences 65.2 (Aug. 1983), pp. 291–307. ISSN: 00255564. DOI: 10.1016/0025-5564(83)90066-4. URL: https://linkinghub.elsevier.com/retrieve/pii/0025556483900664 (visited on 06/12/2023).

[26] Jeng-Huei Chen, Ya-Hui Kuo, and Hsing Paul Luh. “Optimal policies of non-cross-resistant chemotherapy on Goldie and Coldman’s cancer model”. en. In: Mathematical Biosciences 245.2 (Oct. 2013), pp. 282–298. ISSN: 00255564. DOI: 10.1016/j.mbs.2013.07.020. URL: https://linkinghub.elsevier.com/retrieve/pii/S0025556413001843 (visited on 06/12/2023).

[27] Srishti Patil. srishtidoi/two-strike-therapy: Initial Release: Preventing evolutionary rescue in cancer. Version v1.0. Aug. 2024. DOI: 10.5281/zenodo.13332990. URL: 10.5281/zenodo.13332990.

[28] J. B. S. Haldane. “A Mathematical Theory of Natural and Artificial Selection, Part V: Selection and Mutation”. In: Mathematical Proceedings of the Cambridge Philosophical Society 23.7 (1927), pp. 838–844.

[29] H. Allen Orr and Robert L. Unckless. “Population Extinction and the Genetics of Adaptation”. en. In: The American Naturalist 172.2 (Aug. 2008), pp. 160–169. ISSN: 0003-0147, 1537-5323.

[30] Hildegard Uecker, Sarah P. Otto, and Joachim Hermisson. “Evolutionary Rescue in Structured Populations”. en. In: The American Naturalist 183.1 (Jan. 2014), E17–E35. ISSN: 0003-0147, 1537-5323.

[31] Guillaume Martin et al. “The probability of evolutionary rescue: towards a quantitative comparison between theory and evolution experiments”. en. In: Philosophical Transactions of the Royal Society B: Biological Sciences 368.1610 (Jan. 2013), p. 20120088. ISSN: 0962-8436, 1471-2970.

[32] S. P. Otto and M. C. Whitlock. “The Probability of Fixation in Populations of Changing Size”. In: Genetics 146.2 (June 1997), pp. 723–733. ISSN: 0016-6731.

[33] Graham Bell. “Evolutionary Rescue”. In: Annual Review of Ecology, Evolution, and Systematics 48.1 (2017), pp. 605–627.

[34] W.M. Getz. “Optimal control of a birth-and-death process population model”. In: Mathematical Biosciences 23.1 (1975), pp. 87–111. ISSN: 0025-5564. DOI: 10.1016/0025-5564(75)90122-4. URL: https://www.sciencedirect.com/science/article/pii/0025556475901224.

[35] Amaury Lambert. “Probability of fixation under weak selection: A branching process unifying approach”. en. In: Theoretical Population Biology 69.4 (June 2006), pp. 419–441. ISSN: 00405809.

[36] John Lamperti. “Continuous state branching processes”. In: Bulletin of the American Mathematical Society 73.3 (1967), pp. 382–386.

[37] Daniel T. Gillespie. “Exact stochastic simulation of coupled chemical reactions”. In: The Journal of Physical Chemistry 81.25 (1977), pp. 2340–2361. DOI: 10.1021/j100540a008. eprint: https://doi.org/10.1021/j100540a008. URL: https://doi.org/10.1021/j100540a008.

